# The solution structures of higher-order human telomere G-quadruplex multimers

**DOI:** 10.1101/2020.11.13.382036

**Authors:** Robert C. Monsen, Srinivas Chakravarthy, William L. Dean, Jonathan B. Chaires, John O. Trent

## Abstract

Human telomeres contain the repeat DNA sequence 5’(TTAGGG), with duplex regions that are several kilobases long terminating in a 3’ single-stranded overhang. The structure of the single-stranded overhang is not known with certainty, with disparate modes proposed in the literature. We report here the results of an integrated structural biology approach that combines small-angle X-ray scattering, circular dichroism (CD), analytical ultracentrifugation, size-exclusion column chromatography and molecular dynamics simulations that provide the most detailed characterization to date of the structure of the telomeric overhang. We find that the single-stranded sequences 5’(TTAGGG)_n_, with n=8, 12, and 16, fold into multimeric structures containing the maximal number (2, 3, and 4, respectively) of contiguous G4 units with no long gaps between units. The G4 units are a mixture of hybrid-1 and hybrid-2 conformers. In the multimeric structures, G4 units interact, at least transiently, at the interfaces between units to produce distinctive CD signatures. Global fitting of our hydrodynamic and scattering data to a worm-like chain (WLC) model indicates that these multimeric G4 structures are semi-flexible, with a persistence length of about 34 Å. Investigations of its flexibility using MD simulations reveal stacking, unstacking, and coiling movements, which yield unique sites for drug targeting.

## INTRODUCTION

Telomeres are structures found at the end of eukaryotic chromosomes which protect genomic DNA from degradation, end-to-end fusion, and homologous recombination(1,2). The human telomere consists of the repeat d(TTAGGG)_n_, and ranges from 5-25 kb in length with an extended single-stranded 3’ overhang of a few hundred bases in non-germ cells(3). This locus has long been associated with human diseases, such as cancer(4) and telomeropathies(5), as well as aging(6) and general genome homeostasis(7). In normal somatic cells, each round of cellular division results in a shortening of the telomere due to the so-called end replication problem—a mechanism believed to be protective against uncontrolled replication(8). Once the telomere has become critically short in normal (non-stem) cells, a DNA damage response is triggered, resulting in uncapping of the telomere-bound shelterin proteins and, eventually, apoptosis(2,8). Cancer cells avoid this fate by utilizing mechanisms that restore telomere length. In more than 85% of cancers, this is accomplished by reactivating human telomerase reverse transcriptase (hTERT), a ribonucleoprotein that extends the telomere 3’ overhang(9-11). G-quadruplex (G4) formation in the telomere overhang can inhibit hTERT binding and extension function(12). Treating cells with telomere G4-specific small molecules leads to uncapping of the shelterin proteins and a sequestering of the free single-stranded telomere overhang, ultimately resulting in a telomere-specific DNA damage response(13-15). These findings have made telomere G4 an attractive cancer target(15).

G-quadruplexes form in guanine-rich sequences, in which guanine tracts interact to form square planar tetrads (G-tetrads) that stack atop one another and are stabilized by coordinating cations, pi-stacking interactions, and a Hoogsteen hydrogen bonding network(16). Many telomere G4 topologies have been characterized at the atomic level by X-ray crystallography and NMR studies. These studies have demonstrated that the monomeric form of the human telomere can exist as parallel(17), hybrid 3+1(18,19), antiparallel(20), and two-tetrad antiparallel(21) structures under various ionic and crowding conditions. The Yang lab(18,19,22), Patel lab(23-25), and we(26) have since shown that the wild-type telomere adopts primarily the hybrid-1 and hybrid-2 topologies in physiologically relevant solution conditions. The Yang lab has shown by NMR that *in vitro* the wild-type monomeric telomere sequence of the form (TTAGGG)_4_T exists in a dynamic equilibrium of hybrid-2 (~75%) and hybrid-1 (~25%)(27).

Telomere G-quadruplexes have also been observed directly in cells. *In vivo*, G4-specific antibodies and fluorescent ligands have confirmed the formation of telomere G4s(28-30). Using the sequence AGGG(TTAGGG)_3_, Hong-Liang and colleagues used ^19^F-NMR cell studies to show that the hybrid-1, -2, and a two-tetrad anti-parallel type (hybrid-3), but not the parallel or antiparallel “basket” topologies, spontaneously form when injected into live HeLa cells(31). Altogether, these studies demonstrate that the most physiologically and thermodynamically relevant monomeric telomere conformations are of the hybrid type.

Although the monomeric telomere G-quadruplex has been extensively studied, there is little structural information on longer telomere sequences forming higher-order telomere structures. Conservative estimates of the length of the single-stranded overhang of the human telomere in fibroblasts indicate that the sequence exceeds the ~30 nucleotides necessary for formation of a single telomere G-quadruplex. Estimates of “normal” single-stranded overhangs range from ~50 to >600 nucleotides(3,32), supporting the possibility of multiple G4s forming in tandem. There have been few attempts to characterize these systems at the atomic level because of the difficulties involving guanine imino overlap and structural polymorphism which hamper NMR studies(27), and the difficulty of obtaining quality crystals for X-ray diffraction(26). Elucidating this higher-order structure is important, as its role in mediating interactions with shelterin proteins, single-stranded binding proteins, and telomerase is critical in maintaining genomic integrity(1,33,34).

To date, only a few low-resolution molecular models and characterizations were reported for the long telomere sequences. In 2006, using a combination of gel electrophoresis, CD, and UV-melting, Yu *et al*. proposed that the telomere multimer of the form (TTAGGG)_n_, where n is 4, 8, or 12, maximizes its usage of G-tracts by forming a “beads-on-a-string” assembly of a variety of telomere topologies (parallel, antiparallel, and hybrid)(35). In 2009 RenČiuk and colleagues, using CD and PAGE experiments, came to the same general conclusion that the higher-order telomere is capable of folding into multiple conformations in K^+^ buffers (parallel, antiparallel, or hybrid) but also that they have the potential to stack, depending on the amount of macromolecular crowding(36). The same year Xu *et al*. demonstrated the formation of higher-order G-quadruplex formation in 96 nucleotide (nt) long telomere sequences by atomic force microscopy (AFM)(37). While the authors arrived at a similar conclusion about the overall higher order assembly (e.g. maximized G4 formation and potential for G4-G4 interactions), they did not report on the topologies of the G4 subunits. A later AFM investigation of a 96 nt long telomere sequence, (TTAGGG)_16_, by Wang and colleagues reported the presence of gaps between G4 units, and suggested that the extended sequences “rarely” maximize G-tract usage(38). A similar conclusion was drawn from low-resolution studies using electron microscopy (EM), single-molecule magnetic tweezers, and single-molecule force ramp assays(39,40). Although, these studies may suffer from insufficient sample annealing protocols or equilibration times. Our prior biophysical studies investigating the secondary and tertiary structure of the higher-order telomere sequences (TTAGGG)_n_ and (TTAGGG)_n_TT, where n = 4, 8, 12, 16 and 32, gave evidence that these sequences preferentially maximize G-tract usage, and preferentially form a mixture of the hybrid-1 and hybrid-2 conformations(41-43). Subsequent investigations by molecular dynamics (MD) simulations, analytical ultracentrifugation (AUC)(43), and differential scanning calorimetry (DSC)(42) studies indicated that, overall, the extended telomere G4s adopt compact, somewhat rod-like structures via stacking interactions between G4 subunits and intervening TTA linkers(42). The best-fit models from these analyses were alternating (5’) hybrid-1 (3’) hybrid-2, referred to as hybrid-12 and hybrid-121, for n = 8 and n = 12 runs, respectively. Interestingly, thermodynamic studies of these two higher-order systems revealed that “each quadruplex in the higher-order structures is not independent and identical but is thermodynamically unique and is influenced by its neighbors”(42). Clearly, there is no consensus on the higher-order telomere’s behavior in solution. Low-resolution imaging and single-molecule studies would suggest a very flexible beads-on-a-string arrangement with large gaps occurring between G-quadruplexes, whereas the latter investigations suggest a more rigid structure, with maximal G-quadruplex formation.

Using an integrative structural biology approach(44,45), which combines CD, hydrodynamics, molecular dynamics, and small-angle X-ray scattering (SAXS), we show that the telomeric sequences form the maximal number of G4 units without any long gaps. Modeling the hydrodynamic and scattering-derived properties of sequences from 24 nt to 96 nt to a worm-like chain (WLC) model reveals a persistence length of ~34 Å, which is in between that of single-stranded DNA (ssDNA) (~22 Å)(46) and double-stranded DNA (dsDNA) (~550 Å)(47), indicating that the extended telomere G4 is semi-flexible. This flexibility is consistent with MD simulations, which show transient stacking interfaces that create potentially unique binding grooves useful in drug targeting. We follow this with an extensive sequence analysis of the sequence d(TTAGGG)_8_ to determine the major constituent G4 topologies. Using CD and mutational analyses we show that the higher-order human telomere is composed of a ratio of hybrid-1 (~25%) and hybrid-2 (~75%) topologies. Our results are in excellent agreement with prior hydrodynamic and NMR analyses of the human telomere sequences(19,24,43). The resulting structural ensembles provide the first “medium-resolution” look at the conformational heterogeneity and dynamics of the higher-order telomere G-quadruplex.

## MATERIALS AND METHODS

### Oligonucleotides

Oligonucleotide sequences were purchased from IDT (Integrated DNA Technologies, Coralville, IA) with standard desalting. Upon receipt, stock oligos were dissolved in MilliQ ultrapure water (18.2 MΩ x cm at 25°C) at 1 mM and stored at −20.0°C until use. All experiments were carried out in a potassium phosphate buffer (8 mM HPO_4_ ^-2^, 185 mM K^+^, 15 mM Na^+^, 1 mM EDTA^2-^, pH 7.2). Folding was achieved by diluting stock oligos into buffer and boiling in a water bath for 20 minutes, followed by slow cooling overnight. Purification was achieved using size exclusion chromatography (SEC) as detailed previously(48). Briefly, oligos were annealed at concentrations of 40-60 μM, filtered through 0.2 μm filters, and injected onto an equilibrated Superdex 75 16/600 SEC column (GE Healthcare 28-9893-33) using a Waters 600 HPLC system. The flow rate was maintained at 0.5 mL/min and sample fractions were collected every 2 minutes from 100 to 180 minutes run time. The molecular weights of fractionated species were estimated based on a regression analysis of elution time vs. log(MW) of protein standards (Sigma #69385), the major folded species were visually evident as symmetric peaks when monitored at 260 nm (or 280 nm for protein standards). Fractionated samples were pooled and stored at 4°C prior to concentration. Where applicable, pooled fractions were concentrated using Pierce protein concentration devices with 3k MWCO (Thermo #88512, #88515, and #88525) which were rinsed free of glycerol. For AUC and SEC-SAXS experiments, samples were dialyzed after concentration using Spectra/Por Float-A-Lyzers G2 3.5 kDa (Sigma #Z726060) in order to buffer match. Concentrations were determined using molar extinction coefficient given in **Table 1**.

**Table 1.**
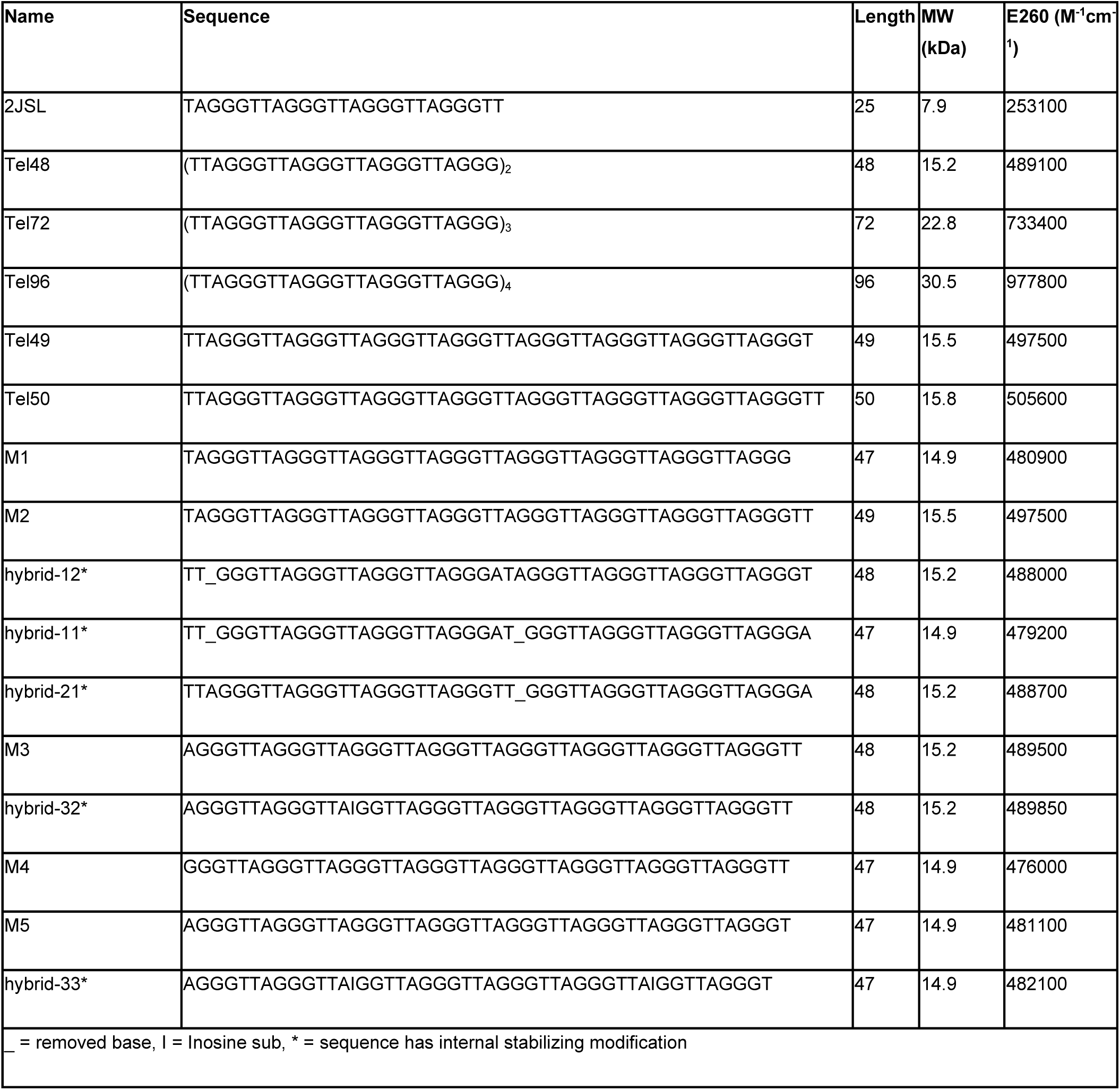
Names, properties, and sequences of oligonucleotides used in this study.

### Size exclusion chromatography (SEC) determination of Stokes radii

Elution times from re-injections of SEC purified fractions were used in the method of Irvine(49) to determine Stokes radii, which were converted to translational diffusion (*D*_*t*_) coefficients for use in Multi-HYDFIT hydrodynamic modeling (see below). Stokes radii were determined from a regression analysis of elution time vs. log(MW) of protein standards (Sigma #69385).

### Analytical ultracentrifugation (AUC)

Sedimentation velocity (SV) experiments were performed in a Beckman Coulter ProteomeLab XL-A analytical ultracentrifuge (Beckman Coulter Inc., Brea, CA) at 20.0°C and 40,000 rpm in standard 2-sector cells using either an An60Ti or An50Ti rotor. Samples were equilibrated in the rotor at 20.0°C for at least 1 hour prior to the collection of 100 scans over an 8-hour period. Initial analyses were performed in SEDFIT(50) using the continuous C(s) model with resolution 100 and S range from 0 to 10. A partial specific volume of 0.55 mL/g for DNA G-quadruplexes was used as previously determined(43). The Tel72 and Tel96 sequence sedimentation coefficients were additionally corrected for any concentration-dependence using three separate concentrations (**Figure S2**).

### Circular dichroism

CD spectra were collected on a Jasco-710 spectropolarimeter (Jasco Inc. Eason, MD) equipped with a Peltier thermostat regulated cell holder equilibrated to 20.0°C. Spectra were collected using the following instrument parameters: 1 cm path length quartz cuvettes, 1.0 nm step size, 200 nm/min scan rate, 1.0 nm bandwidth, 2 second integration time, and 4 scan accumulation. Spectra were corrected by subtracting a buffer blank and normalized to molar circular dichroism (Δε, M^-1^cm^-1^) based on DNA strand concentration using the following equation:

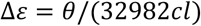

where *θ* is ellipticity in millidegrees, *c* is molar DNA concentration in mol/L, and *l* is the path length of the cell in cm. Comparison or fitting of CD spectra with their monomer theoretical spectra was done manually in Microsoft Excel using spectra from a previously reported database(51). Residual sum of squares (RSS) analysis of the CD ΔΔε “residuals” was carried out and plotted in Origin 2020.

### Size exclusion chromatography resolved small angle X-ray scattering (SEC-SAXS)

SAXS was performed at BioCAT (beamline 18ID at the Advanced Photon Source, Chicago) with in-line size exclusion chromatography. Samples in BPEK buffer (8 mM HPO^4-2^, 185 mM K^+^, 15 mM Na^+^, 1 mM EDTA^2-^, pH 7.2) were loaded onto an equilibrated Superdex 75 10/300 GL column, which was maintained at a constant flow rate of 0.7 mL/min using an AKTA Pure FPLC (GE Healthcare Life Sciences) and the eluate after it passed through the UV monitor was directed through the SAXS flow cell, which consists of a 1 mm ID quartz capillary with 50 μm walls. A co-flowing buffer sheath was used to separate the sample from the capillary walls, helping to prevent radiation damage(52). Scattering intensity was recorded using a Pilatus3 1M (Dectris) detector which was placed 3.5 m from the sample giving access to a q-range of 0.004 Å^-1^ to 0.4 Å^-1^. A series of 0.5 second exposures was acquired every 2 seconds during elution and data was reduced using BioXTAS RAW 1.6.3(53). Buffer blanks were created by averaging regions flanking the elution peak and subtracted from exposures selected from the elution peak to create the I(q) vs. q curves used for subsequent analyses. More information on SAXS data collection, reduction and interpretation can be found in **Table S1**. SAXS sample preparation, analysis, data reduction, and data presentation has been done in close accordance with recent guidelines(54).

### Molecular dynamics simulations and hydrodynamic calculations

Molecular dynamics simulations were carried out on Tel48, Tel72, and Tel96 constructs created previously(41), or modeled based on their solution NMR structures from the Protein Data Bank using the following IDs: 2GKU (hybrid-1)(23), 2JSL (hybrid-2)(24). Base modifications and optimization of starting configurations were performed in UCSF Chimera v1.12(55) or Maestro v11.8(56). The partial negative charges of carbonyls at the center of tetrads were neutralized with coordinated potassium counter-ions added manually in Maestro with subsequent minimization prior to simulation. The PDB structures created were then imported into the xleap module of AMBER 2018(57), neutralized with K^+^ ions, and solvated in a rectangular box of TIP3P water molecules with a 12 Å buffer distance. All simulations were equilibrated using sander at 300 K and 1 atm using the following steps: (1) minimization of water and ions with weak restraints of 10.0 kcal/mol/Å on all nucleic acid residues (2000 cycles of minimization, 500 steepest decent before switching to conjugate gradient) and 10.0 Å cutoff, (2) heating from 0 K to 100 K over 20 ps with 50 kcal/mol/Å restraints on all nucleic acid residues, (3) minimization of the entire system without restraints (2500 cycles, 1000 steepest decent before switching to conjugate gradient) with 10 Å cutoff, (4) heating from 100 K to 300 K over 20 ps with weak restraints of 10.0 kcal/mol/Å on all nucleic acid residues, and (5) equilibration at 1 atm for 100 ps with weak restraints of 10.0 kcal/mol/Å on nucleic acids. The resulting coordinate files from equilibration were then used as input for 100 ns of unrestrained, solvated MD simulations using pmemd with GPU acceleration in the isothermal isobaric ensemble (P = 1 atm, T = 300 K) with DNA OL15 and TIP3P water force fields. Periodic boundary conditions and PME were used. 2.0 fs time steps were used with bonds involving hydrogen frozen using SHAKE (ntc = 2). For the Tel48 constructs, an additional 100 ns of accelerated MD (aMD) simulation were carried out using the average torsional and potential energies from the end of the standard 100 ns simulations as input for calculating the “boosting” of both whole potential and torsional terms (iamd = 3). Trajectories were analyzed using the CPPTRAJ module in the AmberTools18(57) package. Hydrodynamic properties were calculated as average and standard deviation of equally spaced trajectory snapshots (i.e. every 100 ps) using the program HYDROPRO10(58) with the recommended parameters for G-quadruplexes(59). Clustering of the trajectories was performed using the DBSCAN method in the CPPTRAJ module of Amber (minpoints = 5, epsilon = 1.7, sieve 10, rms residues 1-48 over atoms P, O3’, and O5’). Electrostatic calculations for visualization were performed using PDB2PQR software on the APBS web server (http://server.poissonboltzmann.org/)(60,61) with AMBER force field and pH set to 7.2. All molecular visualizations were performed in UCSF Chimera v1.12(55).

### Ensemble optimization method (EOM)

Telomere ensembles were derived using the Ensemble Optimization Method 2.1(62) program from the ATSAS suite of tools. For the Tel48 constructs, which included the four combinations of hybrid-1 and hybrid-2 topologies (i.e. hybrid-11, hybrid-12, hybrid-21, and hybrid-22), a total of 2,000 PDB snapshots were derived from the 100 ns of MD and aMD trajectories stripped of water and K^+^ and pooled, totaling 8,000 coordinate files. GAJOE was used in pool “-p” mode, with maximum curves per ensemble set to 30, minimum curves per ensemble set to 1, constant subtraction allowed, curve repetition allowed, and the genetic algorithm (GA) repeated 200 times. Where noted, the minimum curves were increased to higher numbers, and the curve repetition was disallowed. The same process was repeated for the Tel72 (hybrid-122, -121, -212, and -221) and Tel96 (hybrid-1222, -2122, -2212, -2221) constructs with a total of 4,000 pooled structures. In brief, EOM takes a large pool of macromolecules covering as much conformational space as possible (and reasonable) and selects from this pool a sub-ensemble of conformers that best recapitulate the experimental scattering. The best fitting ensemble is the subset of weighted theoretical curves from conformations that minimizes the discrepancy χ^2^:

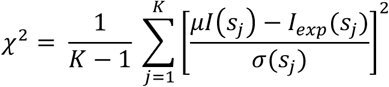

where *I*_*exp*_*(s*_*j*_) is the experimental scattering, *I(s*_*j*_) is the calculated scattering, *K* is the number of experimental points, *σ(s*_*j*_) are standard deviations, and *μ* is a scaling factor(63).

### *Ab initio* model generation and single model validation

P(r) distributions obtained from GNOM(64) using scattering data from 2JSL, Tel48, Tel72, and Tel96 were submitted using the ATSAS online servers (https://www.embl-hamburg.de/biosaxs/atsas-online/) for either *DAMMIN* (2JSL, Tel48) or *DAMMIF* (Tel72, Tel96) bead model generation. Relevant parameters, anisotropy assumptions, normalized spatial discrepancy values (NSDs), *χ*^*2*^ values, and resolutions are given in **Table S1**. Single best fit models for each telomere construct were determined using the initial pool of conformers derived from MD simulations (or NMR structures for 2JSL) and calculated using CRYSOL(65). The best fit structure was determined by minimization of a *χ*^*2*^ function:

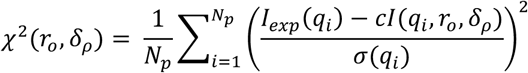

where *I*_*exp*_(*q*_*i*_) and *I*(*q*_*i*_) are the experimental and computed profiles, respectively, *σ(q*_*i*_) is the experimental error of the measured profile, *N*_*p*_ is the number of points in the profile, and *c* is the scaling factor. Two other parameters, *r*_*o*_ and *δ*_*p*_, are fitted and represent the effective atomic radius and the hydration layer density, respectively.

### Flexibility analyses by swollen Gaussian chain and WLC models

Fitting of the radii of gyration, as measured by SEC-SAXS for 2JSL, Tel48, Tel72, and Tel96 was performed as outlined recently by Capp at al.(66) using the following relationship describing the stiffness and conformational space of a swollen Gaussian coil:

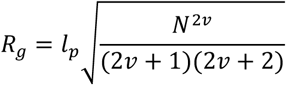

Where *l*_*p*_ is the persistence length and *v*is the Flory coefficient. The *R*_*g*_ values with their respective errors were plotted against their G4 number and fit using a non-linear least squares fitting procedure in Origin 2020 (OriginLab Corporation, Northampton, MA, USA).

For the worm-like chain (WLC) modeling, a global analysis of three properties was used in the program Multi-HYDFIT(67,68). Measurements of two other properties, diffusion coefficient, *D*_*t*_ (calculated from measured Stokes radii, *R*_*s*_, via the Stokes-Einstein equation), and corrected sedimentation coefficient, *S*_*20,w*_, were obtained for each sequence using SEC (average ± S.D. of 4 measurements each) or AUC (concentration series extrapolated to infinite dilution ± standard error from regression analysis), respectively. Each value, with respective weighting and molecular weight (MW), was used as the input for the Multi-HYDFIT program. Multi-HYDFIT uses comparisons of the so-called equivalent radii and ratios of radii to calculate theoretical values of *R*_*g*_, *D*_*t*_, and *S*_*20,w*_, which are then compared to that of the measured values. The ratios of radii are directly related to the ratios of length to diameter (*L/d*) and length to persistence length (*L/l*_*p*_). With starting estimates of *l*_*p*_, *d*, and mass per unit length (*M*_*L*_), the Multi-HYDFIT procedure seeks to minimize a target function(68):

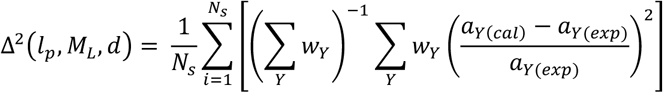

where *N*_*s*_ is the number of samples of different MW, *w*_*Y*_ is the weighting, and *a*_*Y*_ is the ratio of radii for each property. In this equation, the outermost sum runs over the *N*_*s*_ samples and the innermost sum runs over the available properties of each sample. The Δ^2^ is a mean-square relative deviation for the data, and 100Δ is the percent difference between experimental and theoretical values over the entire set. Additional information is required for the calculation, such as temperature (here 20.0°C was used), solvent viscosity (0.00995 poise), starting guesses for diameter, *d* (10 to 100 angstrom), mass per unit length, *M*_*L*_ (10 to 300 Da/angstrom), and persistence length, *l*_*p*_ (20 to 100 angstrom). Intrinsic viscosities calculated from best-fit models using HYDROPRO10 were also included with modest weighting, as they were not empirically determined but rather derived from SAXS best-fit models. The goal of the procedure is to determine the best-fit values of the latter properties, which are given in **Table 2**.

**Table 2.**
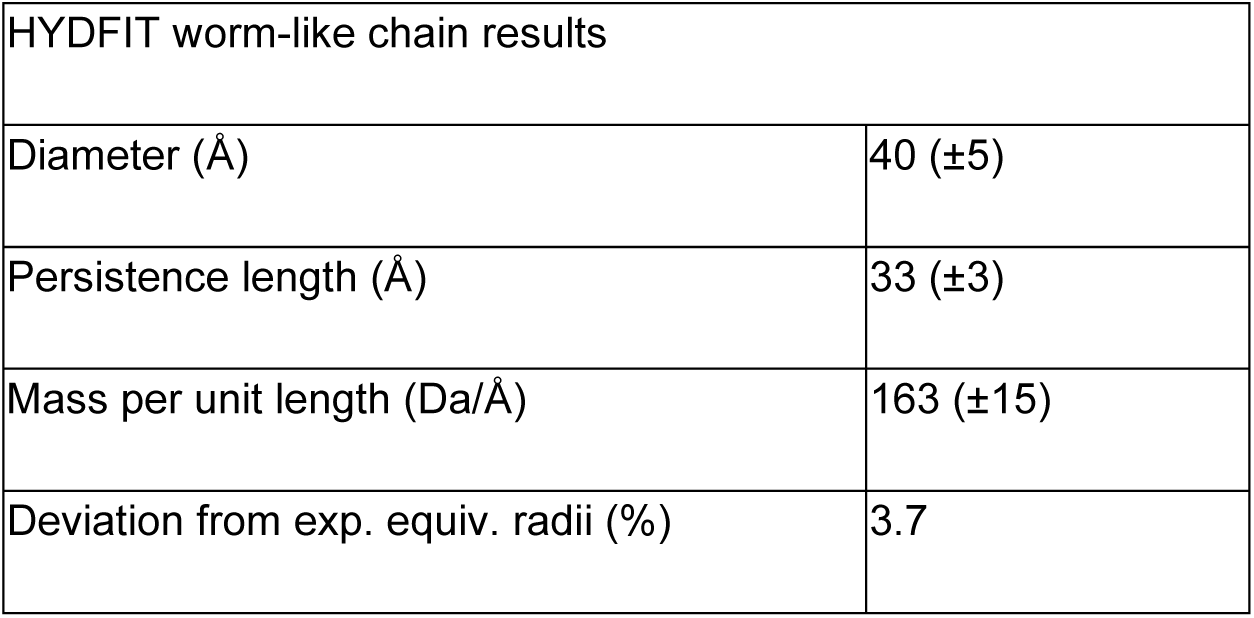
Table of properties derived from Multi-HYDFIT fitting of the higher-order telomere experimental properties to a worm-like chain model.

| HYDFIT worm-like chain results |  |
| --- | --- |
| Diameter ($\text{\AA}$) | 40 ($\pm 5$) |
| Persistence length ($\text{\AA}$) | 33 ($\pm 3$) |
| Mass per unit length ($\text{Da}/\text{\AA}$) | 163 ($\pm 15$) |
| Deviation from exp. equiv. radii (%) | 3.7 |

Force of bending curves generated in **Figure 8** were calculated using the literature persistence length values for single- and double-stranded DNA at cationic conditions similar to used here(46,47), based on the relationship(69):

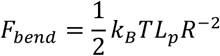

Where *F*_*bend*_ is the bending force in picoNewtons, *K*_*B*_ is the Boltzmann constant, *T* is temperature in Kelvin, *L*_*p*_ is the persistence length in meters, and *R* is the radius of the arc of a curve in meters. The data was plotted such that the values on the X-axis correspond to the end-to-end length of the polymer curved 180° around the arc of a semi-circle.

### Molecular visualizations

All molecular visualizations of MD trajectories and models and RMSD calculations were performed in UCSF Chimera v1.11(55).

## RESULTS

### Small-angle X-ray scattering reveals G4 maximization and indicates that the higher-order telomere G4s are semi-flexible

To verify that the extended telomere sequences are in fact maximizing their G-tract usage we employed size-resolved small-angle X-ray scattering (SEC-SAXS) to assess each sequence for size, shape, and compactness(70,71). The results of the SEC-SAXS analysis for sequences 2JSL (hybrid-2), Tel48, Tel72, and Tel96 (**Table 1**) are shown in **Figures 1, S1**, and **Tables S1. Figure 1A** shows the scattering intensity as a function of momentum transfer (q) on a log-log scale for each sequence. Each scattering profile proceeds horizontally to the Y-axis at low values of q, indicating the absence of inter-particle interactions or repulsions(70). Scattering from 2JSL shows a distinct smooth curvature at higher q values which is indicative of a globular particle, whereas the extended telomere sequences deviate from this curvature between about 0.05 and 0.2 q, suggesting a non-globular structure(71).

**Figure 1.**
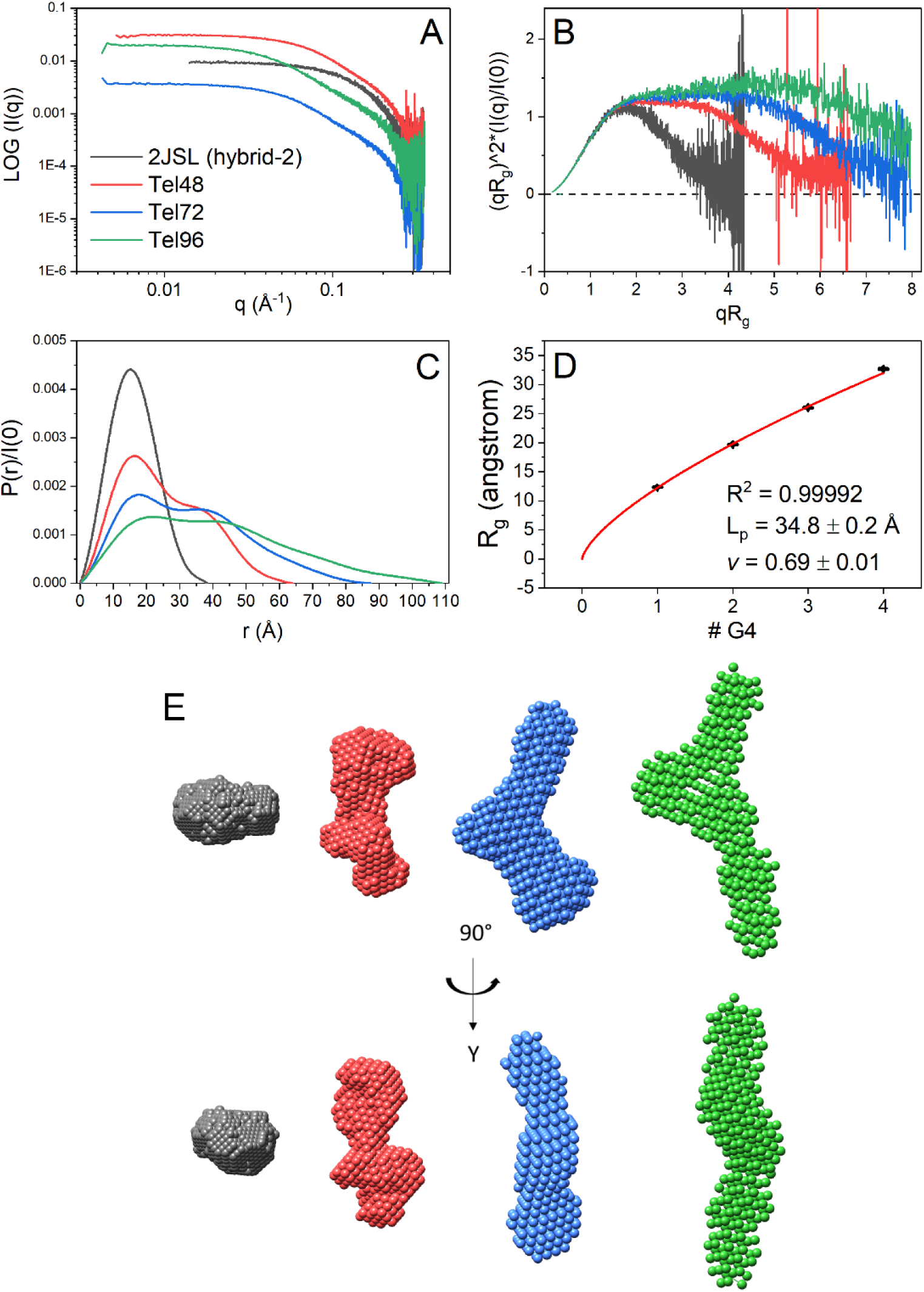
SEC-SAXS analysis of 2JSL (gray), Tel48 (red), Tel72 (blue), and Tel96 (green). (A) Log-log plot of the scattering intensity vs. scattering vector, q. (B) Dimensionless Kratky plots of data in A. (C) Pair distribution function plots of data in A normalized to I(0). (D) Scatter plot of the radii of gyration from each sequence as a function of G-quadruplex motif fit to a swollen Gaussian chain polymer model (see methods) with (inset) derived persistence length (L_p_) and Flory coefficient (*v*). (E) DAMMIN and DAMMIF *ab initio* space-filling models from the data in C.

Two useful transformations of the scattering data are the Kratky plot and distance distribution, P(r), plots (**Figures 1B and 1C**), which allow for qualitative appraisal of compactness and overall structure, respectively(71). In **Figure 1B** the dimensionless Kratky plot shows that 2JSL (gray) exhibits a nearly perfect Gaussian distribution that returns to baseline at high qR_g_, confirming that it is globular and folded(71). The Tel48 and Tel72 sequences also approach baseline at high qR_g_, indicating that they are folded and do not contain significant amounts of flexibility(71). The higher-order sequences also exhibit distinct plateau regions above 2 qR_g_, indicating that they have non-globular shapes and are likely multi-domain, consistent with tandem G4 domains. However, Tel96 (green) exhibits a slight rise in its plateau towards higher qR_g_, indicating that it is flexible relative to 2JSL, Tel48, and Tel72. **Figure 1C** shows the corresponding P(r) distributions (normalized to scattering intensity, I[0]), which are probability distributions of the inner-atomic distances within each macromolecule(70). 2JSL (gray) again exhibits a symmetric distribution, indicative of a globular molecule(71). Conversely, the extended sequences are all multi-modal. Tel48 exhibits a biphasic distribution (red) indicating a characteristic dumbbell-like tertiary arrangement(71), consistent with two G4 domains separated by a small linker region. Tel72 and Tel96 have tri- and tetra-phasic curves, respectively, which we take as indicating three and four contiguous globular domains in tandem, respectively.

P(r) distributions also allow for quantitative characterization of macromolecules. The point on the X-axis at which each sequence converges to zero is the maximum diameter, *D*_*max*_, which is the diameter of the particle’s longest axis(71). The *D*_*max*_ of each sequence increases approximately linearly with a ~24 Å increase with each additional G4 motif. Any substantial amount of telomere species with gaps, or non-maximization of G4s, would likely result in a non-linearity (as well as large upticks in the Kratky curves at high qR_g_). The radius of gyration, *R*_*g*_, is the root mean square distance of the macromolecule’s parts from its center of mass and reflects the particle’s size(70). The *R*_*g*_ can be calculated by either the Guinier approximation (from plots shown in **Figure S1**) or directly from its P(r) distribution, the latter of which is thought to be more representative in cases where flexibility is assumed (although both values should be in general agreement)(71). *D*_*max*_ and *R*_*g*_ values for each sequence are reported in **Table S1**. Shown in **Figure 1D** is a plot of each sequence’s radii of gyration plotted against G4 number. Each additional G4 motif leads to an approximate *R*_*g*_ increase of 6.7 Å.

The extended tails of the P(r) distributions and upward trend in plateau regions of the Kratky plots (**Figure 1B**) signify flexibility. Although plotting the radii of gyration *versus* the putative number of G4 subunits appears entirely linear (R^2^ = 0.9985), a better fit is obtained when fitting to a swollen Gaussian polymer model (R^2^ = 0.9999, **Figure 1D**). The non-linear least-squares fit to this model allows for the estimation of two parameters: persistence length, *L*_*p*_, and the Flory exponent, *v*. The persistence length represents the distance along the telomere G4 polymer which behaves as a rigid rod. At lengths much greater than this, the polymer behaves as a flexible Gaussian chain. The Flory coefficient (also known as the excluded volume parameter) varies between 0.5 and 1.0 and describes the degree of flexibility of the system. A Flory coefficient for a theoretical freely jointed flexible chain is 0.5 (maximum flexibility), whereas that of a rigid rod is 1.0. For reference, the empirical value of chemically denatured proteins is *v* = ~0.588(72). Fitting to this model we find that the telomere G4 has a persistence length of 34.8 ± 0.2 Å and Flory coefficient of 0.69 ± 0.01. The persistence length is approximately the size of a single telomere G4 (~32 Å, calculated from PDB 2JSL less the flanking nucleotides), which indicates that the TTA linkers may provide a point of flexibility. This persistence length is about 50% greater than ssDNA (*L*_*p*_ = ~22 Å under similar ionic conditions(73)). As an independent method of estimating the persistence length, we used the hydrodynamic modeling program Multi-HYDFIT(67). This program integrates multiple independently measured properties, such as sedimentation coefficients (*S*_*20,w*_) from AUC (**Figure S2**)(43), translational diffusion coefficients (*D*_*t*_) from SEC, and radii of gyration (*R*_*g*_) from SEC-SAXS, for a series of macromolecules of given molecular weight (*MW*), and uses these values to find the optimum values of the model parameters for a worm-like chain (WLC) model(68). In total, we fit 12 independent properties from three independent techniques with their respective weights (estimated from standard deviations of multiple measurements), yielding a persistence length of 33 ± 3 Å (**Table 2**), in excellent agreement with the *L*_*p*_ estimated from the swollen Gaussian chain model. Altogether, these results, along with the qualitative information from Kratky and P(r) distributions, suggest that the extended telomere maximizes G4 formation, is closely packed, and is moderately flexible. The flexibility is consistent with rigid G4 units linked by flexible, hinged, interfaces.

### Ab initio and atomistic modeling reveals an ensemble of conformations ranging from entirely stacked and condensed to a coiled “beads-on-a-string” configuration

The above analyses suggest a flexible system which would render the higher-order SAXS data unsuitable for use in *ab initio* bead reconstruction methods. However, upon seeing the resulting space-filling models we were compelled to include them. **Figure 1E** shows the resulting DAMMIN and DAMMIF space-filling models of 2JSL, Tel48, Tel72, and Tel96 created based on the P(r) data in **Figure 1C** (with corresponding fit results tabulated in **Table S1**). Consistent with predictions from the Kratky and P(r) distribution plots Tel48 looks like a dumbbell with two domains roughly the size of the 2JSL reconstruction with a small linker region in the middle. Similarly, Tel72 and Tel96 have what appear to be three and four G-quadruplex domains (indicated by their distinct “bends”), although their resolution is not quite as high as the Tel48 reconstruction (**Table S1**). The similar overall shape and curvature coupled with the flexibility assessment above indicates a non-rod-like structure for telomere sequences with more than two G4 motif repeats. These shapes are generally in accord with previous hydrodynamic investigations based on rigid structures(43), but offer a more detailed and nuanced characterization because the flexibility of the structures can be taken into account.

We next employed an ensemble modeling approach that combined explicit solvent MD-derived models with the ensemble optimization tool GAJOE (of the EOM 2.0 suite)(63). A CD analysis that will follow indicated that the telomere sequences are best represented by a combination of hybrid-1 and hybrid-2 topologies. However, the order in which they occur is not evident, and it may be that the extended sequences are dynamic and interconvert on timescales much longer than is accessible by standard MD simulations (>1 ms). Therefore, we modeled every combination of the simplest multimer system, Tel48. Using the PDB atomic structures for hybrid-1 (PDB ID: 2GKU) and hybrid-2 (PDB ID: 2JSL) we generated each of the four possible combinations: hybrid-11, hybrid-12, hybrid-21, and hybrid-22. Each structure was subjected to 100 ns of both standard MD and accelerated MD (aMD) simulations to produce a pool of 8,000 conformations for use in minimal ensemble and single structure modeling efforts. In the GAJOE ensemble optimization method, a pool of PDB atomic coordinate files are generated that cover as much conformational space as possible and utilized in calculating theoretical scattering profiles. Next, a genetic algorithm acts on these scattering profiles to minimize a fitness function by weighting each scattering profile and comparing combined profiles to the experimental (see methods). The output is an ensemble of conformers which best recapitulate the experimental scattering profile based the minimized *χ*^*2*^ value. An ensemble is considered a better fit than a single conformer when its *χ*^*2*^ value is reduced relative to the single best-fit conformation.

**Figures 2 and S3** shows the results of modeling efforts with the Tel48 constructs. **Figures 2A and 2B** are scatter plots which show the calculated radii of gyration (Y-axis) and corrected sedimentation coefficients (X-axis)(**Figure S2B**), for each of 2,000 frames across both MD (light gray) and aMD (dark gray) trajectories for the hybrid-12, -21, -11, and -22 constructs. These plots indicate that both hybrid-12 and -21 sample conformations which agree with either the experimental *R*_*g*_, *S*_*20,w*_, or intersect both values. The hybrid-11 and hybrid-22 constructs rarely sampled conformations that corresponded with the experimental values (see **Figure S3**). Interestingly, although hybrid-12 extensively samples conformations which agree with both hydrodynamic and scattering-derived measurements, the best fit model by CRYSOL analysis was found to be a highly extended hybrid-21 conformation (cyan dot and curve in **Figures 2B & 2C**). Because this conformation appeared unnatural (e.g. maximally extended) and did not agree very well with the P(r)-derived *R*_*g*_ and *D*_*max*_ values, we speculated that this configuration may be biased simply by our initial start configurations. The hybrid-21 clearly tended towards an overall more compact structure as indicated by the histograms. Therefore, we next asked what the maximum number of curves could be which could reconstruct the experimental scattering without worsening the *χ*^*2*^ value. We found that an ensemble of six conformations gave approximately the same *χ*^*2*^ value (magenta dots in **Figures 2A** and **2B**, magenta curve **Figure 2C**) and agreed much better with the experimental *R*_*g*_ and *D*_*max*_ values from the P(r) analysis (*R*_*g,cal*_ *=* 19.58 Å *vs. R*_*g,exp*_ = 19.69 Å and *D*_*max,calc*_ = 66 Å *vs. D*_*max,exp*_ = 65 Å, **Table S1**). The resulting topologies were a 50/50 mix of hybrid-12 and -21 (**Figure 2D**), which sampled conformations ranging from extended to fully stacked. The flexibility of the ensemble was only marginally lower than the pool, as judged by EOM’s Rflex flexibility analysis, supporting semi-flexibility. Interestingly, we found that one of the hybrid-12 conformers (bottom right of **Figure 2D**) was nearly identical in conformation to our previously reported hybrid-12 model(41), with an RMSD of just 1.6 Å over all residue pairs (**Figure S4**).

**Figure 2.**
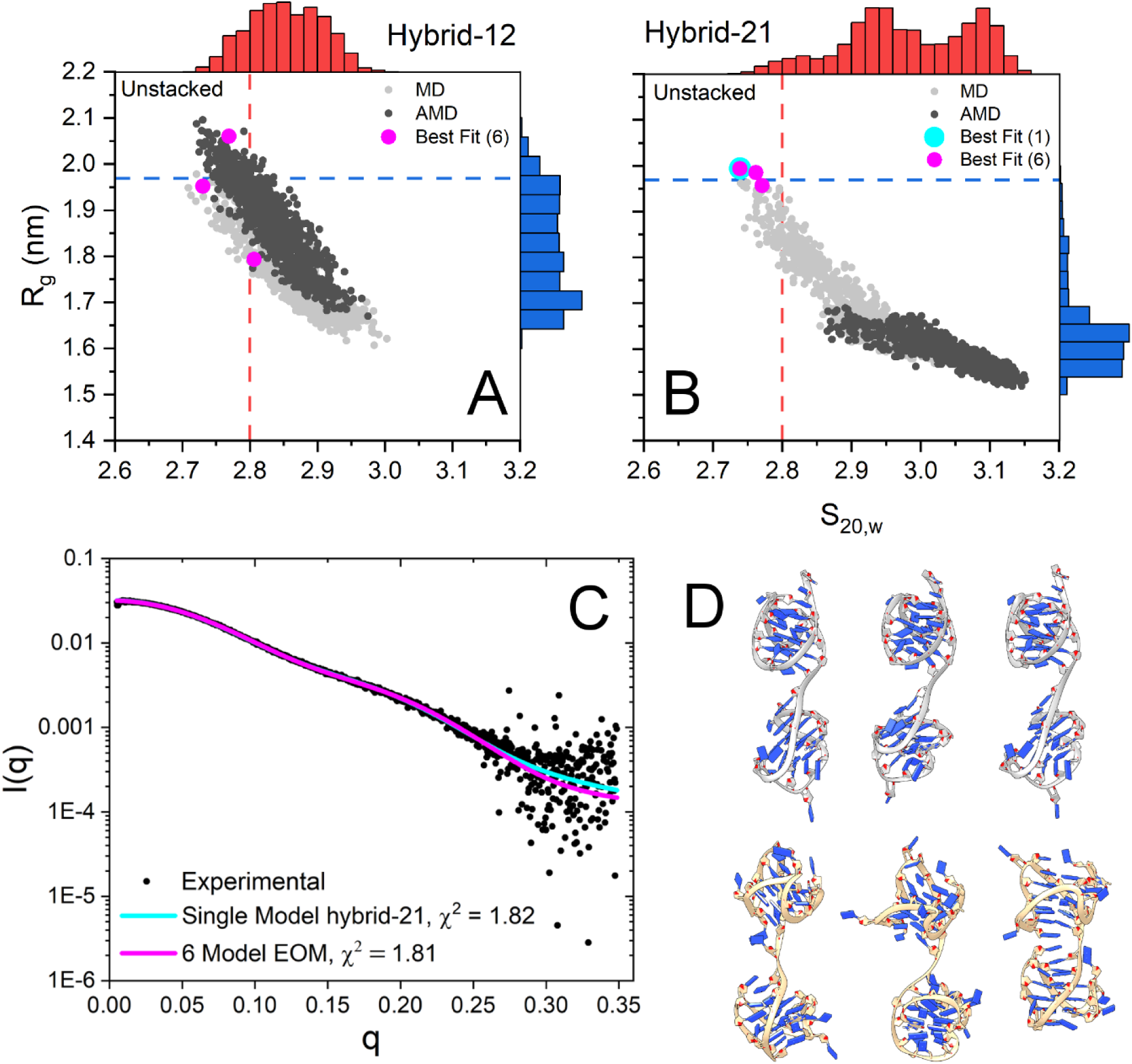
Results of Tel48 SAXS atomistic modeling efforts. (A-B) scatter plots of calculated radii of gyration and sedimentation coefficients for hybrid-12 (A) and hybrid-21 (B) with MD-derived values shown in light gray and aMD-derived values in dark gray. The inset dashed red and blue lines represent the experimentally measured values for sedimentation coefficient and radius of gyration, respectively. The outer histograms represent the distributions of values from both MD and aMD snapshots combined. The cyan dot represents the single best-fit model (hybrid-21) as determined by CRYSOL (top left model in D). Magenta dots represent the six conformers in the best fit ensemble (all six models in D). (C) Experimental SAXS scattering data with fits from single (cyan) or ensemble (magenta) calculated scattering overlaid with χ^2^ values inset. (D) Single best fit model (hybrid-21, top left model) and best fit ensemble of six conformers (top row hybrid-21, bottom row hybrid-12). Models are oriented with their 5’ ends at the top.

Next, we investigated the Tel72 and Tel96 constructs in the same manner as above but only using standard MD. Models were created to reflect the ratio of hybrid-1/-2 (25/72) as determined by our later CD analyses of telomere mutants (e.g. hybrid-121, -122, -212, and -221 for Tel72 and hybrid-1222, -2122, - 2212, and -2221 for Tel96). The hybrid-121 was included because it was proposed previously(41). Each model was then subjected to explicit solvent MD and simulated for a total of 100 ns. From these trajectories, 1,000 equally spaced frames were used as the pool for EOM’s GAJOE program. For the Tel72 the best fit was obtained with an ensemble of four conformers (*X*^*2*^_*ensemble*_ = 1.09 *vs. X*^*2*^_*single-model*_ = 1.81) which was composed of the hybrid-212 (89.7%) and hybrid-221 (10.3%) (**Figure 3**). Surprisingly, this configuration agrees well with Tel48 having a 5’ preference for hybrid-2 followed by hybrid-1. The *R*_*g*_ and *D*_*max*_ values of the Tel72 ensemble (**Figure 3C**) agree with the experimental values (*R*_*g,calc*_ = 25.8 Å *vs. R*_*g,exp*_ = 26.0 Å and *D*_*max,calc*_ = 83 Å *vs. D*_*max,exp*_ = 87 Å), indicating that the ensemble is an excellent solution. Similarly, Tel96 scattering was best recapitulated by an ensemble of four conformers (*X*^*2*^_*ensemble*_ = 1.15 *vs. X*^*2*^_*single-model*_ = 2.08) but was composed entirely of different conformations of the hybrid-2122 (**Figure S5**). Again, there is an agreement with a 5’ hybrid-2 followed by hybrid-1. The *R*_*g*_ and *D*_*max*_ values of this conformer ensemble are also in agreement with the experimental values (*R*_*g,calc*_ = 32.1 Å *vs. R*_*g,exp*_ = 32.7 Å and *D*_*max,calc*_ = 103 Å *vs. D*_*max,exp*_ = 109 Å), indicating that this ensemble is reasonable. In both cases, the EOM Rflex quantification of flexibility indicates that the ensembles are only marginally less flexible than the pool (**Table S1**), consistent with the system semi-flexibility. This flexibility is also illustrated by the conformer ensembles themselves (**Figures 3C and S5**). Further, docking of each ensemble into their respective *ab initio* space-filling models from **Figure 1E** reveals excellent fits for the models of topological sequence 5’-hybrid-2,-1,-2,-2 (**Figure 4**). Collectively, these analyses indicate that in physiological buffer conditions the extended telomeres maximize their formation of G4 subunits, prioritize hybrid-2 at the 5’ end, and are semi-flexible.

**Figure 3.**
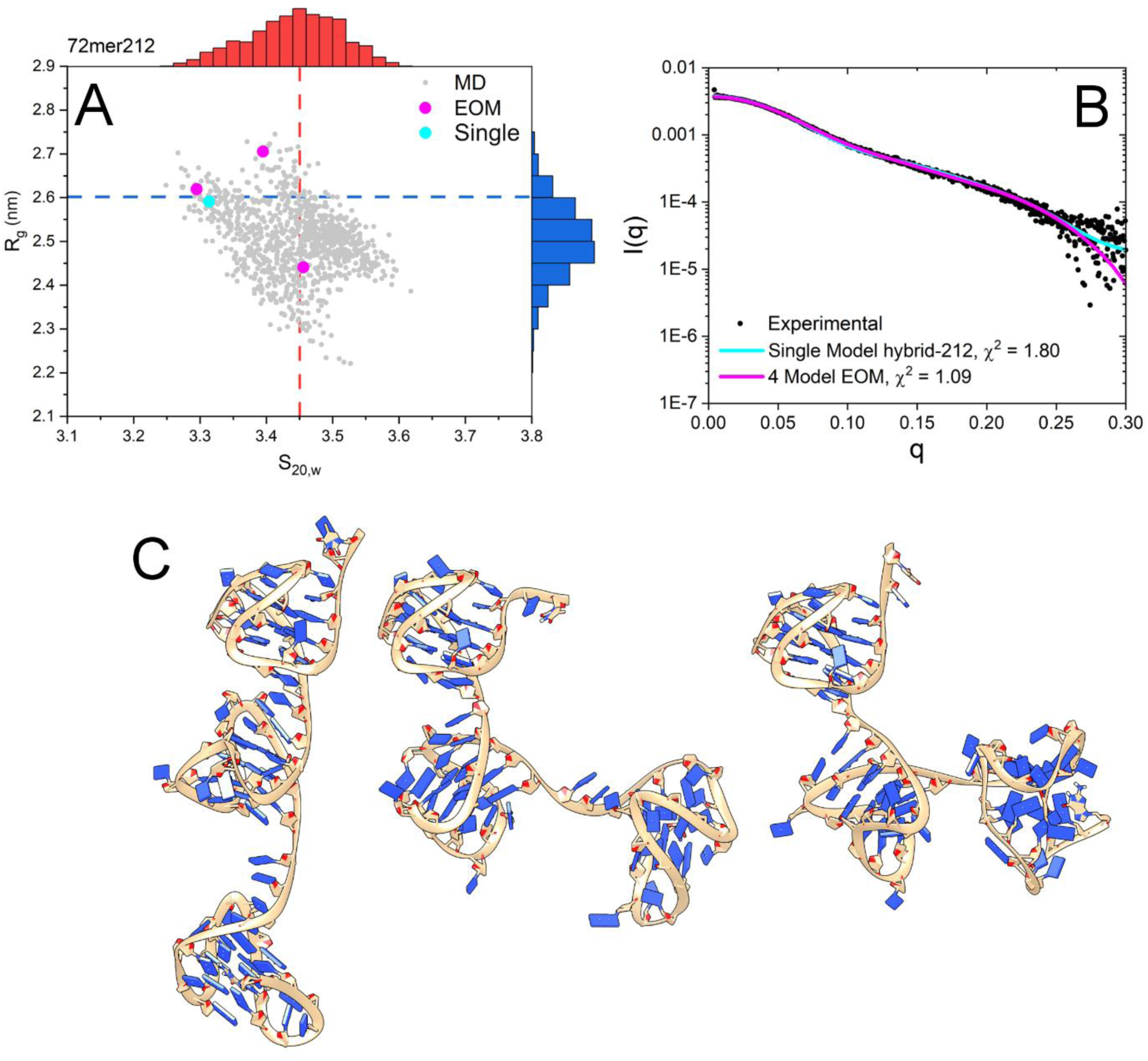
Results of Tel72 SAXS atomistic modeling efforts. (A) scatter plot of calculated radii of gyration and sedimentation coefficients for the hybrid-212 model from 100 ns of standard MD simulation. The inset dashed red and blue lines represent the experimentally measured values for sedimentation coefficient and radius of gyration, respectively. The outer histograms represent the distributions of values. The cyan dot represents the single best-fit model as determined by CRYSOL. Magenta dots represent the four conformers in the best fit ensemble. (B) Experimental SAXS scattering data with fits from single (cyan) or ensemble (magenta) calculated scattering overlaid with χ^2^ values inset. (C) Conformations of the three hybrid-212 configurations (not showing the hybrid-221) from the best fit ensemble as determined by EOM. Models are oriented with their 5’ ends at the top.

**Figure 4.**
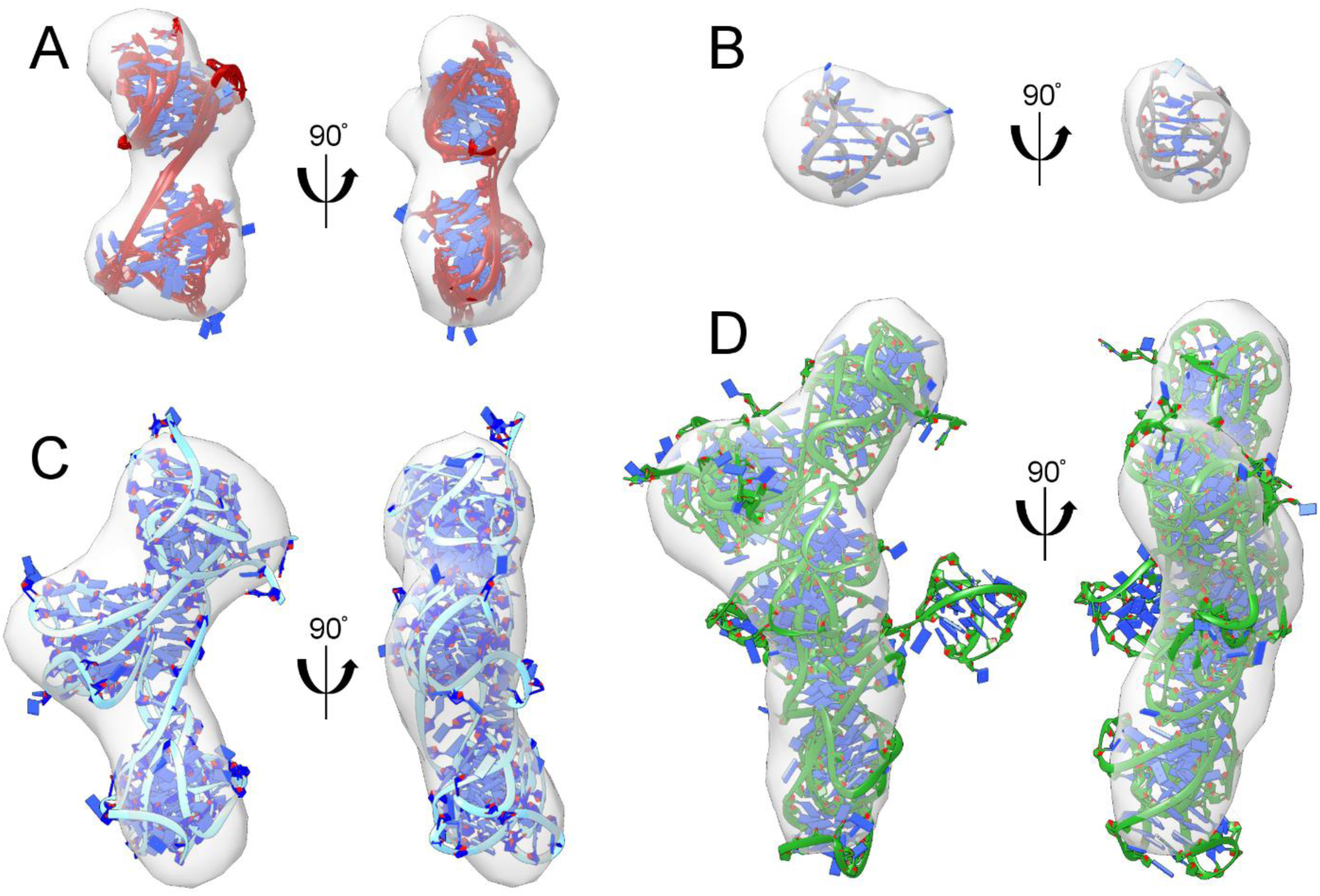
Telomere G4 ensembles from EOM GAJOE analysis docked into *ab initio* space-filling reconstructions from DAMMIN/DAMMIF. (A) Tel48 hybrid-21 conformers, (B) 2JSL with single best-fit NMR-derived model, (C) Tel72 hybrid-212 conformers (the same as in **Figure 3C**), (D) Tel96 hybrid-2122 models (the same as in **Figure S5**).

### The Tel72 hybrid-212 preferentially samples a stacked conformation, forming unique electronegative binding pockets useful for drug targeting

The Tel72 ensembles reflect well the SAXS-derived properties, *R*_*g*_ and *D*_*max*_. However, they do not agree as well with their measured sedimentation coefficients. SAXS scattering is exquisitely sensitive to changes in particle volume (conformation in this case). Systems which exist in an equilibrium of stacked and unstacked, or coiled and beads-on-a-string, will have a scattering profile which is composed of a continuous distribution of conformations (as the scattering intensity, I[0], is directly related to the volume of the scattering particle)(74). Therefore, we next endeavored to find the most frequently sampled conformation from the MD trajectory of the Tel72 hybrid-212 construct. **Figure 5A** shows the top three most frequently sampled conformations across the 100 ns simulations with their respective weighting (% of MD frames). This analysis suggests that approximately 47% of the frames from simulation sampled a configuration which was partially (middle) or entirely stacked (left and right models). The major stacked conformation sampled by hybrid-212 has a calculated sedimentation coefficient which is in excellent agreement with the experimental (*S*_*20,w(calc)*_ = 3.45 *versus S*_*20,w(exp)*_ = 3.46, **Figure S2**) although the calculated radius of gyration is slightly lower (*R*_*g(calc)*_ *=* 2.45 nm *versus R*_*g(exp)*_ *=* 2.60). Electrostatic calculations of the most prominent form reveal highly electronegative grooves, which are appropriately sized for small molecules (**Figure 5B**). Overall, these analyses show that the hybrid-212 model of the Tel72 is consistent with all available spectroscopic, hydrodynamic, X-ray scattering, and MD-based analyses, and forms unique junctional grooves for selective small molecule targeting.

**Figure 5.**
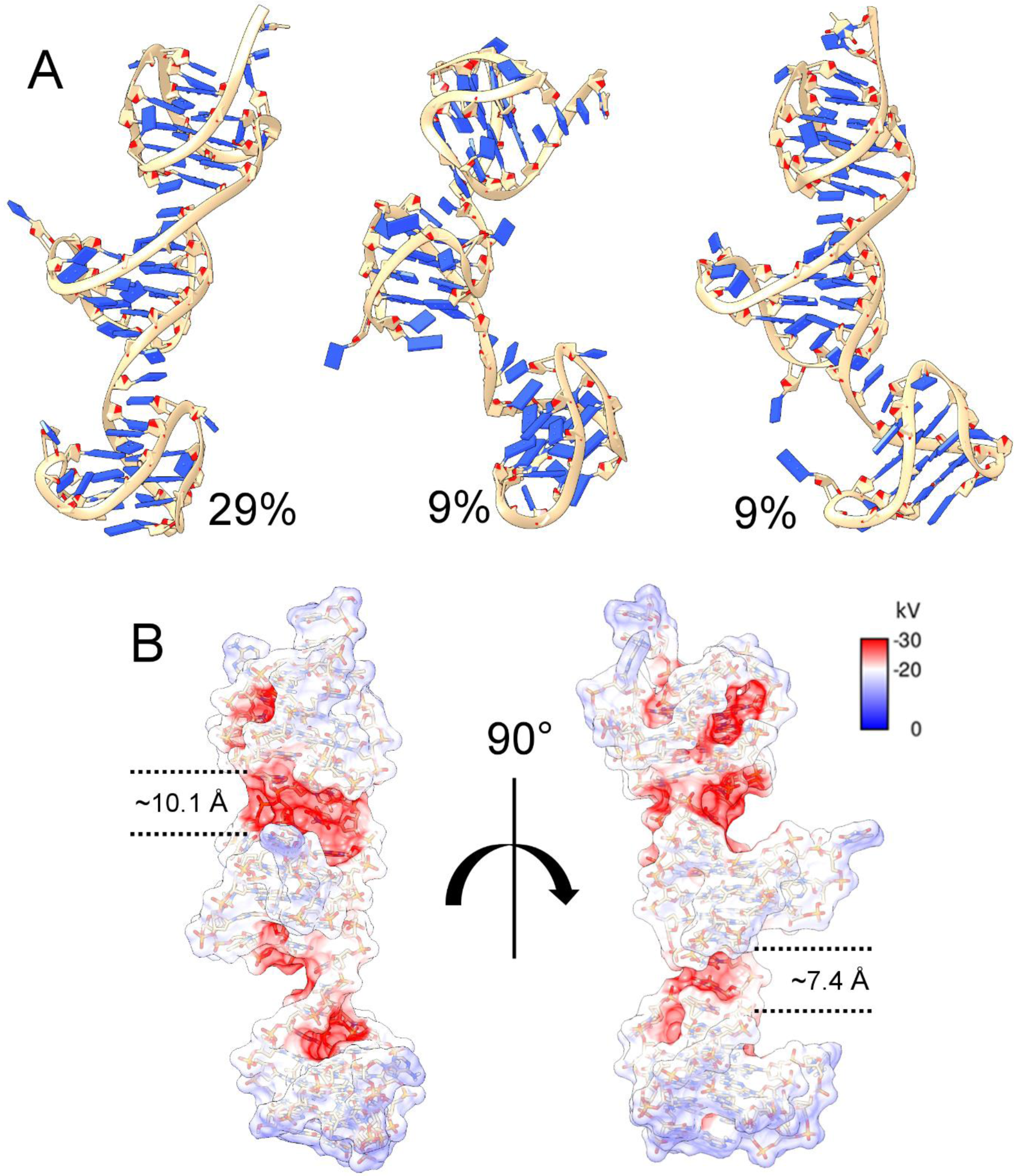
Results of MD clustering analysis of the Tel72 hybrid-212. (A) Top three representative centroids of DBSCAN clusters accounting for ~47% of frames across the entire 100 ns trajectory. (B) space-fill electrostatic APBS map of the first model from A with dashed lines indicating the approximate sizes of each groove created at the two stacking interfaces.

### The major topologies of the extended telomere G4 are hybrid-1 and hybrid-2

Simultaneous with our structural investigations above, we investigated the conformations of G4 units within higher-order structures by using systematic sequence variations of the wild type (WT) Tel48 sequence and observing changes in circular dichroism spectra. These sequence “mutants” were created with variation in terminal nucleotides or by changes in internal sequences that favor the various hybrid topologies (e.g. hybrid-1 and hybrid-3) (**Table 1**). The Tel48 spectrum (black line in **Figure 6A**) has a main peak at ~290 nm, pronounced shoulder from 265-275 nm, and a trough at 235 nm, indicating that it is primarily composed of hybrid type folds(27). Comparing sequence variants of the form T_n_AGGG(TTAGGG)_7_T_m_, where *n* = 1 or 2, and *m* = 0, 1, or 2, we found no major spectral differences (**Figure S6A**), indicating that changes in these flanking nucleotides have no effect on the overall topology. Removal of the 5’-end thymine residues led to a modest reduction in the shoulder at ~270 nm and peak at 290 nm when compared to the WT sequences of similar length (**Figure S6B**, red and blue lines). We speculated that this could be due to the formation of hybrid-3 in the 5’-most G4 unit, which has been reported in shorter sequences lacking the 5’ thymine residues(21,31). Indeed, when an inosine is introduced to favor the hybrid-3 topology in the 5’-most putative G4 the CD changes observed at 270 and 290 nm become more pronounced (**Figure S6C**, blue solid line). Importantly, this suggests that the hybrid-3 topology is not a major topology, as the extended telomere (in the cell) will always include 5’ thymine residues. The spectral change due to hybrid-3 incorporation is made more evident when stabilized in both 5’ and 3’ G4 units (**Figure S6C**, purple), which is of the same shape but approximately 2x the magnitude of hybrid-3. Thus, the hybrid-3 is not likely to exist in the context of the extended telomere, aside from as a potential folding intermediate(21).

**Figure 6.**
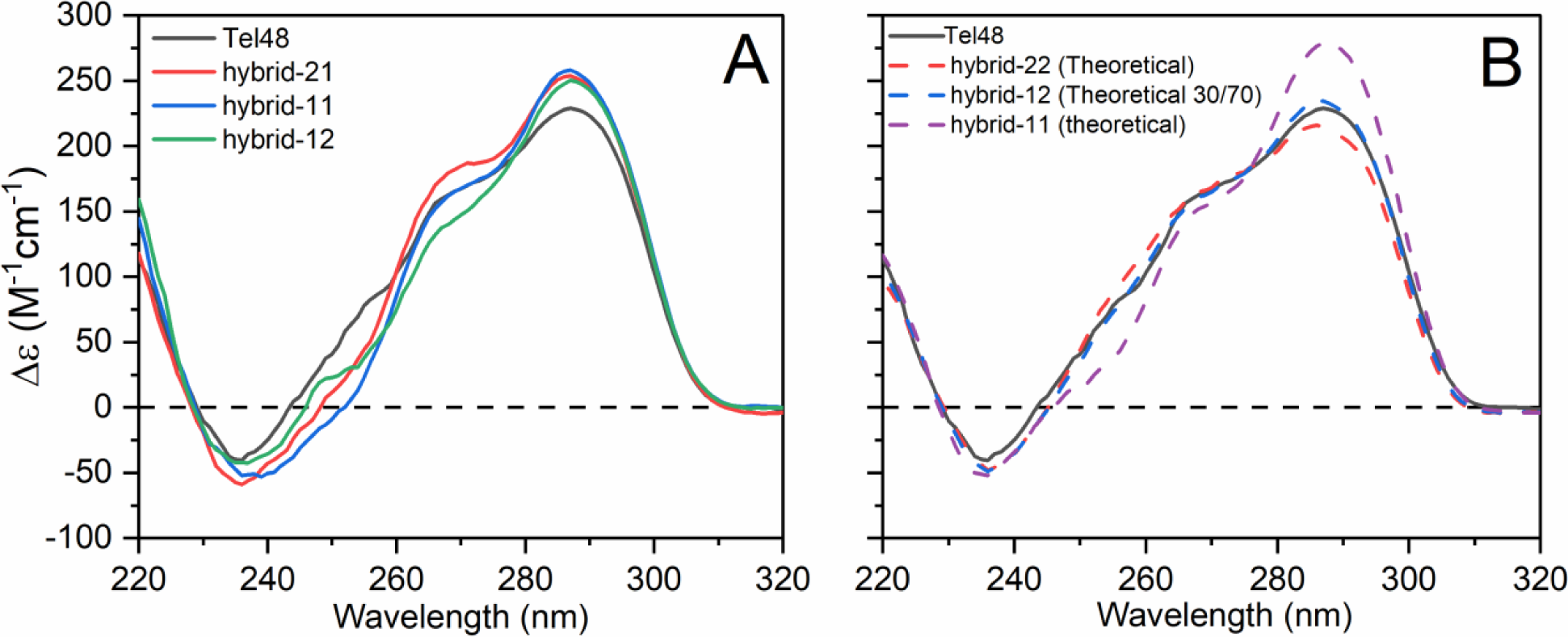
Normalized circular dichroism spectra of Tel48 mutants and theoretical monomer G4 spectra. (A) CD spectra comparison of the WT Tel48 G-quadruplex (black) with constructs created to favor the hybrid-1 form in the second (red), first and second (blue), or first position (green). (B) Comparison of the Tel48 CD spectrum with theoretical monomer CD combinations of hybrid-22 (red dashed), hybrid-12 with a 30/70 weighting (blue dashed), and a hybrid-11 (purple dashed).

Comparison of the hybrid-21, -11, and -12 sequences to Tel48 revealed subtle differences in each case (where hybrid-2 is assumed as the major conformation in unadulterated telomere sequence flanked by thymine at both ends) (**Figure 6A**). Overall, each spectrum was consistent in shape, but varied in magnitude at various wavelengths. We suspected that this may indicate a preference for the hybrid-22 form. A theoretical hybrid-22 spectrum overlaid nicely with Tel48 (**Figure 6B**, red dashed line). In contrast, a theoretical hybrid-11 (**Figure 6B**, purple dashed line) had a greatly increased 290 nm peak and slight reduction at ~250 nm and looked similar to the mutant hybrid-11 spectrum from **Figure 6A**. Based on the reported 25/75 ratio of hybrid-1 and hybrid-2 for the monomer telomere sequence flanked at both ends by thymine(75), we next tested a variety of computed weighted combinations of hybrid-1 and -2 spectra and found that a 30/70 ratio best reflects the Tel48 spectrum (compare blue dashed line with black in **Figure 6B**). Collectively, these results indicate a preference for both hybrid-1 and -2 topologies in the extended telomere sequence, consistent with our EOM analysis of Tel48.

### CD indicates that the higher-order telomere sequences converge on a 25/75 ratio of hybrid-1 and hybrid-2 with maximization of G4 formation

Strand-normalized circular dichroism spectra are the sum of constituent secondary and tertiary structure(76). Thus, just as above we expect that the spectra of higher-order telomere G4s could be recreated by addition of their measured “monomer” spectra. However, comparisons of the various monomer spectra (hybrid-1,-2,-3, and basket forms) to our higher-order telomere spectra resulted in non-negligible “difference” spectra of roughly the magnitude expected for di- or tri-nucleotides. As prior studies suggest, and we have shown here, the extended telomere sequences maximize their G4 potential by leaving no G-tract gaps. The resulting stacking junctions, or other inter-G4 interactions that constrain the loop regions, could potentially give rise to a “junctional” CD signal(76). Thus, to generate the theoretical “junctional” spectrum, we utilized the Tel48 mutant spectra from above and subtracted from them theoretical constituent monomer spectra as appropriate. The resulting spectrum is shown in **Figure 7A**. The junction spectrum has a peak at 240 nm and troughs at ~260 and ~285 nm, which is consistent with the known CD spectra of adenine and thymine polynucleotides(77). This spectrum was then used as a correction factor, and was subtracted from the Tel48, Tel72, and Tel96 spectra (**Figure 7B**). A plot of Δε_290_ *versus* the putative number of G4s yields a linear regression with a near zero Y-intercept, which is more physically relevant than the regression without the correction (**Figure 7C**). The slope of the corrected regression data indicates that each additional putative G4 increases Δε_290_ by ~128 M^-1^cm^-1^, in excellent agreement with the average Δε_290_ obtained from hybrid (3+1) monomers. These corrected spectra should now be a composite spectrum of monomer G-quadruplex components. A novel finding here is that the CD spectra of higher-order G4 structures contain discernable contributions from G4-G4 interactions.

**Figure 7.**
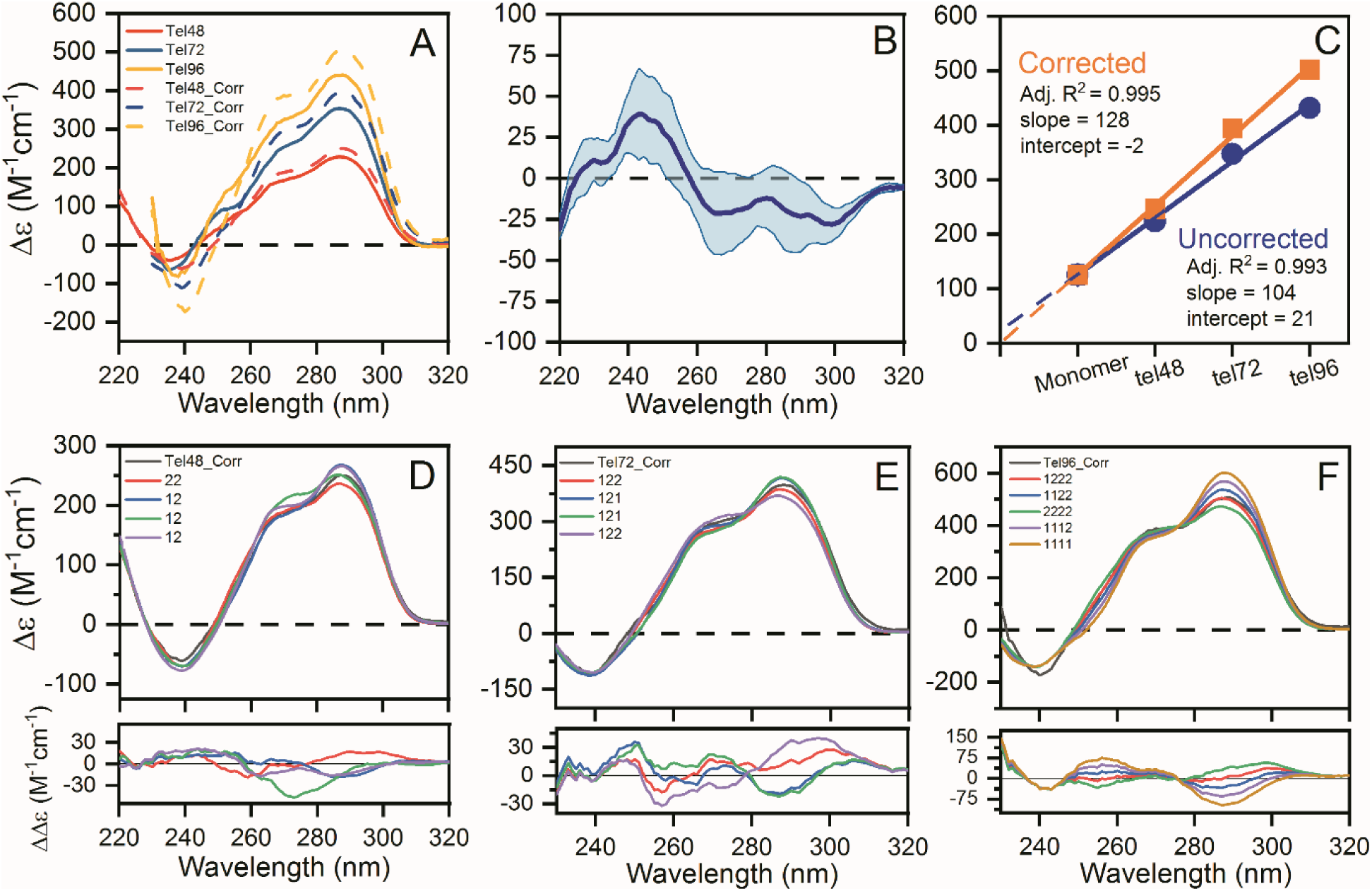
Circular dichroism analysis of the higher-order telomere G-quadruplexes. (A) Pre- and Post-corrected (“Corr”) CD spectra of the Tel48, Tel72, and Tel96 sequences by subtraction of the “junctional” spectrum in B. (B) The average (dark blue line) and range (light blue space fill) of “junctional” CD spectra derived from deconstruction of the Tel48 sequences in **Figures 6 and S6** using constituent monomer spectra. (C) Regression analysis of the uncorrected and corrected Δε_290nm_ values as a function of the number of G4 motifs. (D-F) Corrected CD spectra of the Tel48, Tel72, and Tel96 sequences with overlaid theoretical spectra derived from the linear addition of monomer spectra. The red spectrum in each plot is the best fit as judged by RSS analysis. Residuals are shown below each figure. See **Figure S7** for the full RSS analysis along with the PDB identifiers of sequences used to compute theoretical spectra.

The hybrid-1, -2, and -3, as well as the antiparallel basket monomer spectra were systematically compared with the Tel48, Tel72, and Tel96 corrected spectra. **Figures 7D-F** show the corrected spectra in black with “_Corr” indicating the corrected spectrum. Shown below are residuals from the best fit combinations of monomer spectra. In each plot the red spectrum is the best fit, followed by blue, etc. based on residual sum of squares (RSS) analysis (**Figure S7**). The optimal fit to Tel48_Corr is hybrid-22 (in agreement with **Figure 6A**), followed by various hybrid-1/-2 combinations; Tel72_Corr is best fit by a combination of hybrid-1, -2, and -2; Tel96 is best fit by a hybrid-1, -2, -2, -2 (not necessarily in that order as shown above). See **Figure S7** for the exhaustive residual sum of squares (RSS) ordering, PDB IDs, and distributions of CD residuals for each fit. These results suggest an overwhelming preference for hybrid-2 in the longer sequences, although some other hybrid-1/-2 combinations also yield comparable fits. Altogether, the above analyses confirm that the primary two topologies making up the WT telomere higher-order G4s are hybrid-1 and hybrid-2, with proportions approaching a 25% hybrid-1, 75% hybrid-2. Further, the linearity and slope of the regression analysis and excellent agreement with theoretical fits indicates that no long gaps exist in the higher-order human telomere.

## DISCUSSION

These results provide the most detailed characterization of extended human telomere G-quadruplex structures in solution to date. We combine circular dichroism, hydrodynamics, and small-angle X-ray scattering experiments integrated with available high-resolution NMR studies on monomeric G4 structures(27) to build medium resolution higher-order structures. For dimeric structures containing two contiguous G4 units (Tel48), the best model is one featuring a 5’ hybrid-2, followed by hybrid-1. For longer sequences with three and four G4 units, a mixture of hybrid-2 and hybrid-1 conformations seems to be present in an approximate 3:1 ratio. Our results show unequivocally that for all sequences up to 96 nt in length, the human telomere sequence maximizes its G4 formation, leaving no gaps—in direct contrast to prior EM, AFM, and single-molecule force ramping investigations(38-40). Our results provide the first quantitative estimates of the rigidity of folded telomeric DNA through determination of its persistence length (*L*_*p*_ *=* ~34 Å). We find that the semi-flexibility of the telomere G-quadruplex is best modeled by an ensemble of configurations which fluctuate between an entirely stacked multimer and unstacked monomeric G4s, as observed by MD simulations, providing potentially unique sites for small molecule targeting in the junctional regions. This model suggests that rigid G4 units are connected by a short, dynamic, interfacial hinge. That interface constitutes a unique structural element to target in drug design efforts.

The WT human telomere monomer sequences exhibit a high degree of polymorphism *in vitro*(27). Under physiologically relevant K^+^ solution conditions the WT telomere G4 adopts a hybrid type conformation, favoring hybrid-2 over hybrid-1 (19,22). This conformational bias is seemingly dictated by the presence of 5’ or 3’ flanking nucleotides. Addition of 5’-TTA to the core sequence, GGG(TTAGGG)3, leads to the favoring of hybrid-1 by a 5’-end capping adenine triplet, whereas addition of one or two thymine to the 3’-end results in a favoring of the hybrid-2 form via a T:A:T triplet stack on the 3’-end(27). The extended, end-flanked sequence, (TTAGGG)4T, forms a major configuration of hybrid-2 (~75%), with minor amounts of hybrid-1 (~25%)(75). This implies that the energy barrier between the two forms may be small. Consistent with this, our mutational analysis by CD and modeling studies agree with a 75/25 ratio of hybrid-2/-1 for the higher-order WT species. A significant result from our higher-order CD analyses is the unique junctional spectrum (**Figure 7B**). This spectrum was useful in providing a rationale for why the higher-order species exhibited CD signatures that were lower than expected for maximized G4 formation. Moreover, both SAXS and MD modeling studies revealed favorable, but dynamic, stacking interfaces between G4 moieties, in agreement with thermodynamic analyses(42).

Prior NMR investigations of the WT telomere sequence, (TTAGGG)4T, indicate a dynamic equilibrium of conformations(27). If a similar equilibrium exists in the higher-order telomere, then an ensemble of both tertiary conformation and secondary structure would be required to explain both CD and SAXS results. Consistent with this, the Tel48 scattering is modeled well by an ensemble with a 50/50 ratio of hybrid-12 and hybrid-21 conformers. The single model hybrid-21 fit is comparable to the ensemble, and so this solution is, overall, somewhat ambiguous; although, the lack of thymine residues at the 3’-end would, in theory, favor hybrid-1 in the second position, giving us confidence in a preferential hybrid-21 model. Our previous investigation of the WT Tel50 sequence(41) (which differs in sequence by two additional thymine residues at the 3’ end) proposed that the major form is hybrid-12. We used steady-state fluorescence measurements of 2-aminopurine-substitutions to assess the solvent accessibility for each adenine site. From this it was found that residue A15 is the least solvent exposed, which agreed with SASA calculations of the hybrid-12 model (in this conformation A15 is buried in the stacking junction between the two G-quadruplex units). Coincidentally, by having the EOM algorithm increase the number of conformers in the Tel48 ensemble, we find that part of the new solution is a hybrid-12 conformer that is almost identical to the previously proposed Tel50 hybrid-12 model (**Figure S4**). Thus, the collective experimental observables support an equilibrium of hybrid-1 and hybrid-2 in either position.

We have also investigated the possibility of a hybrid-3 form, which is a two-tetrad antiparallel G-quadruplex that has been characterized *in vitro* in potassium containing solution(21), and confirmed as existing in a cellular environment by Bao et al.(31). In both instances the telomere sequences used were lacking 5’ thymine residues. The 5’ thymine residue destabilizes the hybrid-3 structure and, ultimately, favors the hybrid-1(21). In the cell the 5’-flanking thymine is always present. We have confirmed that the extended WT telomere sequences do not favor the hybrid-3 in solution by mutational analysis, showing that it may only occur in the 5’-most position when thymine is removed (**Figure S6**). Thus, the hybrid-3 topology may only function as a folding intermediate(21), rather than as a major constituent topology of the higher-order telomere G4.

The single-stranded telomere overhang is a critical regulator of genomic integrity. Spanning the junction of the duplex and single-stranded telomere region is a protective protein complex known as shelterin(1,78). Shelterin is composed of the proteins TRF1, TRF2, RAP1, TIN2, TPP1, and POT1. POT1 (protection of telomeres 1) is essential in sequestering the free 3’ overhang, shielding it from eliciting aberrant single-stranded DNA damage responses, preventing homologous recombination, and regulating the activity of telomerase(34). POT1 binds directly to the 3’ single-stranded overhang with high affinity and in a highly sequence specific manner(1,33,79). EM micrographs have revealed that POT1-TPP1 complexes coat the entirety of the extended telomere 3’ overhang, forming compact, ordered complexes of ssDNA-POT1-TPP1 without gaps(80). Importantly, disruption of POT1’s shielding of the single-stranded overhang elicits an ATR-dependent DNA damage response through the promiscuous ssDNA binding protein RPA(34,81). A recent AFM investigation of the Tel96 sequence with POT1 by the Opresko lab(38) found that maximization of G4 formation “rarely” occurs, and that POT1 associates by simply recognizing the resulting gaps. We, and others(35,36,43,82), find this conclusion at odds with solution-based results. Accessible ssDNA gaps between G4s would allow RPA to compete unimpeded with POT1 binding. Indeed, RPA outcompetes POT1-TPP1 binding to single-stranded telomere DNA *in vitro*(83). Further, G-quadruplex secondary structure enhances POT1-TPP1’s protection against RPA in physiologically relevant levels of K^+^ (150 mM)(84). We recently showed that POT1 unfolds and binds to telomere G4s using a conformational selection mechanism(82) and demonstrated that the kinetics of unfolding the Tel48 sequence is essentially the same as the Tel24 monomeric G4. Importantly, this suggests that a maximization of G4 formation does not impede POT1 binding. Taken together, the physiological significance of telomeric G4 maximization is two-fold: (1) G-quadruplex secondary structure prevents promiscuous RPA binding and, (2) the G4 secondary structure promotes exclusive interaction with shelterin through specific POT1 unfolding and binding, tilting the scale in favor of POT1 over RPA.

The mechanistic details of how the shelterin complex orchestrates the sequestration of the single-stranded 3’ end are still not entirely understood, but are of great importance in drug discovery (78,85). Recently, the Cech laboratory conducted a thorough investigation of co-expressed and isolated complexes of the shelterin proteins *in vitro*(78). Based on their results a shelterin “load & search” model was proposed, whereby TRF2 and POT1 localize the shelterin complex to the telomere by specifically recognizing and binding to the single-stranded/double-stranded (ss/ds) telomere junction. The authors propose that POT1 searches for its optimal binding sequence, TTAG, at the 3’ end by a scanning search mechanism, eventually looping the 3’ end back forming a loop bridged by shelterin proteins that is “unlike a T-loop”. An earlier report from the Cech laboratory found that POT1 and POT1-TPP1 completely coat the long ssDNA telomere repeats *in vitro*(80). Thus, the “normal” sequestration mechanism of the human telomere 3’ overhang is seemingly a POT1-coated loop structure anchored to the ss/ds telomere junction.

DNA looping is a common theme in the cell(69). From a physical stand-point, DNA looping has been extensively studied for its relationship with genetic packaging into nucleosomes and effects on transcription(69,86). A commonly reported measure of the structural rigidity of a biological polymer is the persistence length, *Lp*, that defines the length over which a polymer remains unbent in solution. In this work, we have applied both SAXS and hydrodynamic modeling methods to derive an estimate of the telomere G4 persistence length. Using our telomere G4 *Lp* estimate, along with values reported for single- and double-stranded DNA, we can compare the relative forces required to bend each 180° around the arc of a semi-circle of a given radius (**Figure 8**)(69). From this plot we find that, in the case of single-stranded telomere DNA of length >200 angstroms (~63 nt) the force required to bend the polymer 180° (in the shape of a semi-circle) is negligible—energy requirements on the order of thermal fluctuations. However, if that same stretch of 63 nucleotides were to form maximal G-quadruplexes (decreasing polymer length to <100 angstrom), the increase in energy to bend increases by an order of magnitude—now requiring energy input equivalent to ATP hydrolysis. Although somewhat intuitive, this implies that in the absence of significant energy input, short telomere G4s must be made single-stranded in order to bridge the terminal 3’ TTAG repeat (capped by POT1) with the shelterin complex. More importantly, this figure indicates that small molecules which bind the inter-G4 grooves, thereby increasing its effective persistence length, could shift the force curve to the right (towards the dsDNA curve, red) and subsequently drive up the energy cost for associating the POT1-bound 3’ end with the shelterin loop. Indeed, during the drafting of this manuscript Gao *et al*. have demonstrated that a small molecule targeting the wild-type Tel48 can shift the distribution of conformations to favor a more compact, likely stacked, conformation(87)—a transition that would directly affect persistence length. It is well established that telomere G-quadruplex interacting small molecules are able to displace shelterin components, uncap the telomeres, and ultimately, inhibit telomerase *in vitro* and *in vivo*(14,88). Thus, there is abundant rationale for future work targeting these unique G4 junctional sites with stabilizing small molecules.

**Figure 8.**
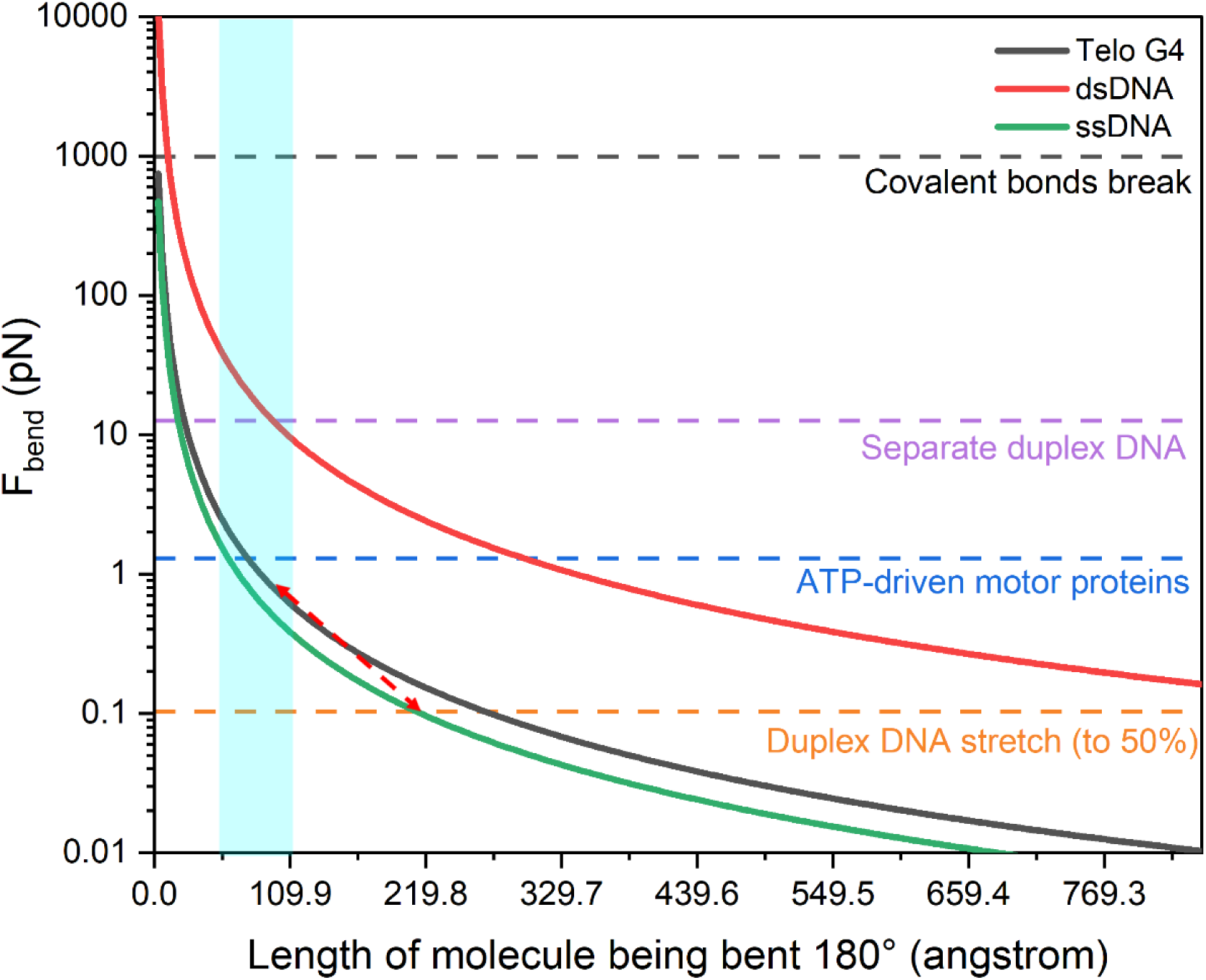
DNA force of bending plot for single-stranded (green), double-stranded (red), and G4 telomere DNA (black). Force curve calculations were performed similar to reference(69) using literature values of persistence length for ssDNA (*L*_*p*_ = 22 Å), dsDNA (*L*_*p*_ = 550 Å), and Telomere G4 (*L*_*p*_ = 34.8 Å) as measured here. The Y-axis is the estimated force (in pN) to bend a length of DNA (X-axis) 180° about the arc of a semi-circle (i.e. if you have a 330 angstrom long single-stranded DNA it will require a force of ~0.05 pN to bend it into a semi-circle). Dashed horizontal lines are visual references to common biological forces found in the cell (orange indicates the approximate range of force from thermal fluctuations). The light blue region highlights the range in which short telomere G4s would be found, indicating that a large force would be required to bend short telomeres (≤96 nt). The dashed red arrow illustrates that if a ~200 Å long ssDNA telomere (approximately 63 nt) were to spontaneously fold into a contiguous G4 structure, the resulting bending force required for a 180° turn increases by an order of magnitude. The increase in bending force is comparable to the same length of DNA in duplex form (330 Å long duplex requires external forces equivalent to ATP hydrolysis to bend 180°). In the case of duplex DNA, the energy requirement of “tight” bending is usually compensated for by the highly positive charge on histones.

Utilizing a robust integrative approach, we have presented here the highest-resolution view of the higher-order telomere G4 to date. SAXS refinement of MD-derived models constructed from high-resolution techniques is now a mainstay in structural biology. However, SAXS refinement of MD generated atomistic models, while excellent for discarding unrealistic topologies and conformations, is not necessarily definitive when conformational and topological polymorphism presents itself. Thus, we await higher-resolution techniques that can inform on the distributions of topologies in the higher-order telomere G-quadruplex.

## DATA AVAILABILITY

Small-angle X-ray scattering data has been deposited in the publicly accessible Small Angle Scattering Biological Data Bank (https://www.sasbdb.org/) under the IDs: SASDKF3 (2JSL), SASDKG3 (Tel48), SASDKH3 (Tel72), and SASDKJ3 (Tel96).

## ACKNOWLEDGEMENTS

This research used resources of the Advanced Photon Source, a U.S. Department of Energy (DOE) Office of Science User Facility operated for the DOE Office of Science by Argonne National Laboratory under Contract No. DE-AC02-06CH11357. This project was supported by grant 9 P41 GM103622 from the National Institute of General Medical Sciences of the National Institutes of Health. Use of the Pilatus 3 1M detector was provided by grant 1S10OD018090-01 from NIGMS.

The content is solely the responsibility of the authors and does not necessarily reflect the official views of the National Institute of general Medical Sciences or the National Institutes of Health.

Molecular graphics and analyses performed with UCSF Chimera, developed by the Resource for Biocomputing, Visualization, and Informatics at the University of California, San Francisco, with support from NIH P41-GM103311

