## Supplemental Material for "The solution structures of higher-order human telomere G-quadruplex multimers"

### TABLE AND FIGURE LEGENDS

**Table S1.** Tabulated collection parameters, data reduction methods, and data analyses for small-angle X-ray scattering data.

**Figure S1.** Guinier analyses of 2JSL, Tel48, Tel72, and Tel96 (left) with fit overlaid in yellow for each sequence and (right) residuals of fits. Guinier fit results are tabulated in Table S1.

**Figure S2.** Sedimentation velocity analysis of higher-order telomere sequences. (Top) Analysis of Tel72 and Tel96 concentration dependence on sedimentation. Extrapolation to the Y-axis gives the infinite dilution $S_{20,w}$ values. (Bottom) representative C(s) vs. $S_{20,w}$ distributions for 2JSL, Tel48, Tel72, and Tel96.

**Figure S3.** Additional results of Tel48 SAXS atomistic modeling efforts shown in **Figure 2** of main text. (A-B) scatter plots of calculated radii of gyration and sedimentation coefficients for hybrid-11 (A) and hybrid-22 (B) with MD-derived values shown in light gray and aMD-derived values in dark gray. The inset dashed red and blue lines represent the experimentally measured values for sedimentation coefficient and radius of gyration, respectively. The outer histograms represent the distributions of values from both MD and aMD snapshots combined. The histograms indicate that the major sampled conformations in both cases are much more compact than would be expected from either SAXS or AUC analyses.

**Figure S4.** Comparison of the Tel48 (cyan) and Tel50 (tan) hybrid-12 conformers. The tan hybrid-12 conformer was taken from an earlier report by Petraccone et al.(1) and the cyan hybrid-12 is the model derived here from EOM (bottom right-most conformer in **Figure 2D**). The pair-wise residue RSMD is 1.6 Å as determined by the matchmaker module of UCSF Chimera v1.12. Potassium is shown as purple spheres and is derived from the Petraccone model (EOM hybrid-12 potassium is hidden).

**Figure S5.** Results of Tel96 atomistic modeling efforts. Depicted are the four hybrid-2122 conformers derived from EOM analysis of the 100 ns MD simulations with their respective weights (as % of the reconstructed scattering curve). All conformers are arranged with their 5' ends at the top of the figure.

**Figure S6.** Normalized CD spectra of Tel48 compared to various flanking residue and internal mutant sequences.

**Figure S7.** Violin plots of residuals obtained as the difference between experimental and theoretical CD reconstruction curves for Tel48, Tel72, and Tel96. The residuals are plotted from best (left) to worst (right) based on residual sum of squares analysis in Origin 2020.

(a) Sample Details.

| | 2JSL | Tel48 | Tel72 | Tel96 |
|---|---|---|---|---|
| Organism | synthetic | synthetic | synthetic | synthetic |
| Source | IDT | IDT | IDT | IDT |
| Extinction coefficient (nearest neighbor approximation) ($M^{-1}$ $cm^{-1}$) | 253100 | 489000 | 733400 | 974870 |
| vbar ($cm^3/g$) (estimate) | 0.55 | 0.55 | 0.55 | 0.55 |
| M from chemical composition (Da) | 7879 | 15212 | 22849 | 30486 |
| SEC-SAXS column, 10 x 300 Superdex 75 | | | | |
| Loading concentration (mg/mL) | 7.0 | 13.0 | 10.0 | 6.0 |
| Injection volume (µL) | 300 | 300 | 250 | 440 |
| Flow rate (mL/min) | 0.7 | 0.7 | 0.7 | 0.7 |
| Solvent (solvent blanks taken from SEC flow through prior to elution of protein) | 8 mM $PO_4^{2-}$, 185 mM KCl, 1 mM EDTA, pH 7.2 | 8 mM $PO_4^{2-}$, 185 mM KCl, 1 mM EDTA, pH 7.2 | 8 mM $PO_4^{2-}$, 185 mM KCl, 1 mM EDTA, pH 7.2 | 8 mM $PO_4^{2-}$, 185 mM KCl, 1 mM EDTA, pH 7.2 |

(b) SAXS data-collection parameters.

| | |
|---|---|
| Instrument/data processing | BioCAT facility at the Advanced Photon Source beamline 18ID with Pilatus3 1M (Dectris) detector |
| Wavelength (Å) | 1.033 |
| Beam size (µm) | 150 (h) x 25 (v) |
| Camera length (m) | 3.5 |
| q measurement range ($Å^{-1}$) | 0.004-0.4 |
| Absolute scaling method | N/A |
| Normalization | To incident intensity, by ion chamber counter |
| Monitoring for radiation damage | Automated frame-by-frame comparison of relevant regions |
| Exposure time, number of exposures | 0.5 s exposure time with a 2s total exposure period (0.5 s on, 1.5 s off) of entire SEC elution |

| | |
|---|---|
| Sample configuration | SEC-SAXS. Size separation by an AKTA Pure with a Superdex 75 Increase 10/300 GL column. SAXS data measured in a 1.5 mm ID quartz capillary |
| Sample temperature (°C) | 20 |

| (c) Software employed for SAXS data reduction, analysis, and interpretation. | |
|---|---|
| SAXS data reduction | Radial averaging; frame comparison, averaging, and subtraction done using BioXTAS RAW 1.6.3 (Hopkins et al. 2017(2)) |
| Extinction coefficient estimate | Nearest neighbor approximation |
| Basic analyses: Guinier, P(r), $V_p$ | Guinier fit, Kratky analysis, and molecular weight using BioXTAS RAW 1.6.3, P(r) function using PRIMUSqt (ATSAS v2.8.4(3)) |
| Shape/bead modelling | DAMMIF (Franke & Svergun, 2009) via ATSAS online (https://www.embl-hamburg.de/biosaxs/atsas-online/) |
| Atomic structure modelling | CRYSOL from PRIMUSqt in ATSAS v2.8.4(Svergun et al., 1995(4)) |
| Three-dimensional graphic model representations | UCSF Chimera v1.11 |

(d) Structural parameters.

| Guinier analysis | 2JSL | Tel48 | Tel72 | Tel96 |
|---|---|---|---|---|
| I(0) ($cm^{-1}$) | 0.00957 ± 0.00002 | 0.0318 ± 0.00002 | 0.003742 ± 0.000005 | 0.0202 ± 0.0000406 |
| $R_g$ (Å) | 12.41 ± 0.05 | 19.23 ± 0.03 | 25.37 ± 0.06 | 31.68 ± 0.13 |
| qmin ($Å^{-1}$) | 0.014 | 0.009 | 0.007 | 0.007 |
| $qR_g$ max | 1.27 | 1.33 | 1.32 | 1.17 |
| Coefficient of correlation, $R^2$ | 0.972 | 0.998 | 0.996 | 0.996 |
| M from volume of correlation, $V_c$ (ratio to predicted) | 6600 (0.84) | 16100 (1.06) | 23500 (1.03) | 30900 (1.01) |
| P(r) analysis (GNOM) | | | | |
| I(0) ($cm^{-1}$) | 0.00954 ± 0.00002 | 0.032 ± 0.00002 | 0.00376 ± 0.000005 | 0.0203 ± 0.00004 |
| Rg (Å) | 12.33 ± 0.03 | 19.69 ± 0.03 | 26.01 ± 0.06 | 32.65 ± 0.10 |
| Dmax (Å) | 38 | 65 | 87 | 109 |
| $\chi^2$ | 0.95 | 1.76 | 0.84 | 1.1 |

| | | | | |
|---|---|---|---|---|
| Porod volume (Å$^{-3}$) (ratio $V_p$/calculated M) | 9040 ( 1.15) | 16100 (1.06) | 26300 (1.15) | 32700 (1.07) |

(e) Shape model-fitting results

| | 2JSL | Tel48 | Tel72 | Tel96 |
|---|---|---|---|---|
| Ambimeter (default parameters) | | | | |
| Number of compatible shape categories, ambiguity score | 19, 1.279 | 634, 2.802 | 712, 2.852 | 644, 2.809 |
| 3D reconstruction | potentially unique | highly ambiguous | highly ambiguous | highly ambiguous |
| DAMMIF (default parameters, 20 calculations) | | | | |
| q range for fitting (Å$^{-1}$) | - | - | 0.0088-0.3146 | 0.0054-0.2539 |
| Symmetry, anisotropy assumptions | - | - | P1, prolate | P1, prolate |
| NSD (standard deviation), No. of clusters | - | - | 1.204 (0.082), 4 | 1.126 (0.095), 11 |
| $\chi^2$ | - | - | 1.166 | 1.167 |
| Resolution (from SASRES) (Å) | | | 31 ± 3 | 38 ± 3 |
| DAMMIN (default, slow) | | | | |
| q range for fitting (Å$^{-1}$) | 0.014-0.3488 | 0.0115-0.3488 | - | - |
| Symmetry, anisotropy assumptions | P1, none | P1, prolate | - | - |
| $\chi^2$, CORMAP P-values | 1.508, 0.1322 | 1.713, 0.249 | - | - |
| Constant adjustment to intensities | 8.46E-05 | 0.00E+00 | - | - |

(f) Atomistic modelling.

| | | | | |
|---|---|---|---|---|
| Crystal structures/atomic coordinate files | PDB ID: 2JSL | Modeled | Modeled | Modeled |
| q range for modelling | 0.01-0.3 | 0.006-0.3 | 0.007-0.3 | 0.0054-0.2500 |
| EOM GAJOE 2.1 (min ensembles = 1, max = 20, default parameters) | | | | |
| $\chi^2$ | - | 1.81 | 1.09 | 1.154 |

| | | | | |
|---|---|---|---|---|
| Rflex (random) / Rsigma | - | 79.26 (88.83) / 0.62 | 79.96 (85.17) / 0.97 | 79.80 (86.06) / 1.39 |
| Constant subtraction | - | 0 | 0 | 0 |
| No. of representative structures | - | 6 | 4 | 4 |
| Final ensemble Rg (Å), Dmax (Å) | - | 19.58, 65.62 | 25.78, 82.65 | 32.11, 103.18 |
| CRYSOL (single model, default parameters) | | | | |
| $\chi^2$ | 1.20 | 1.82 | 1.81 | 2.08 |
| Predicted Rg (Å) | 12.3 | 19.82 | 25.74 | 32.63 |
| Dro (optimal hydration shell contrast), Ra (optimal atomic group radius (Å) | 0.060, 1.760 | 0.065, 1.800 | 0.045, 1.400 | 0.075,1.400 |
| (g) SASBDB IDs for data and models. | | | | |
| ID | SASDKF3 | SASDKG3 | SASDKH3 | SASDKJ3 |

**Table S1.** Tabulated collection parameters, data reduction methods, and data analyses for small-angle X-ray scattering data.

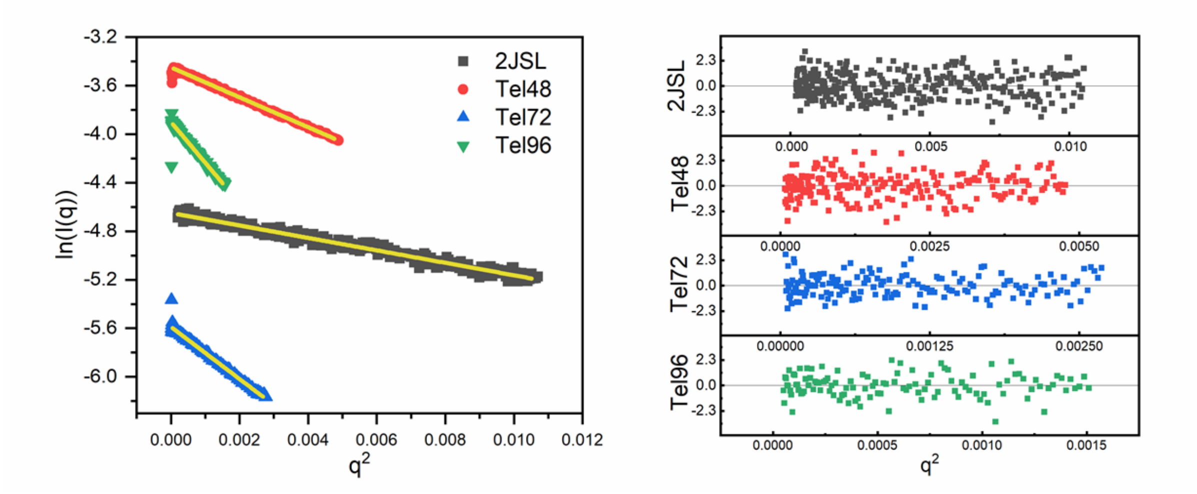

**Figure S1.** Guinier analyses of 2JSL, Tel48, Tel72, and Tel96 (left) with fit overlaid in yellow for each sequence and (right) residuals of fits. Guinier fit results are tabulated in Table S1.

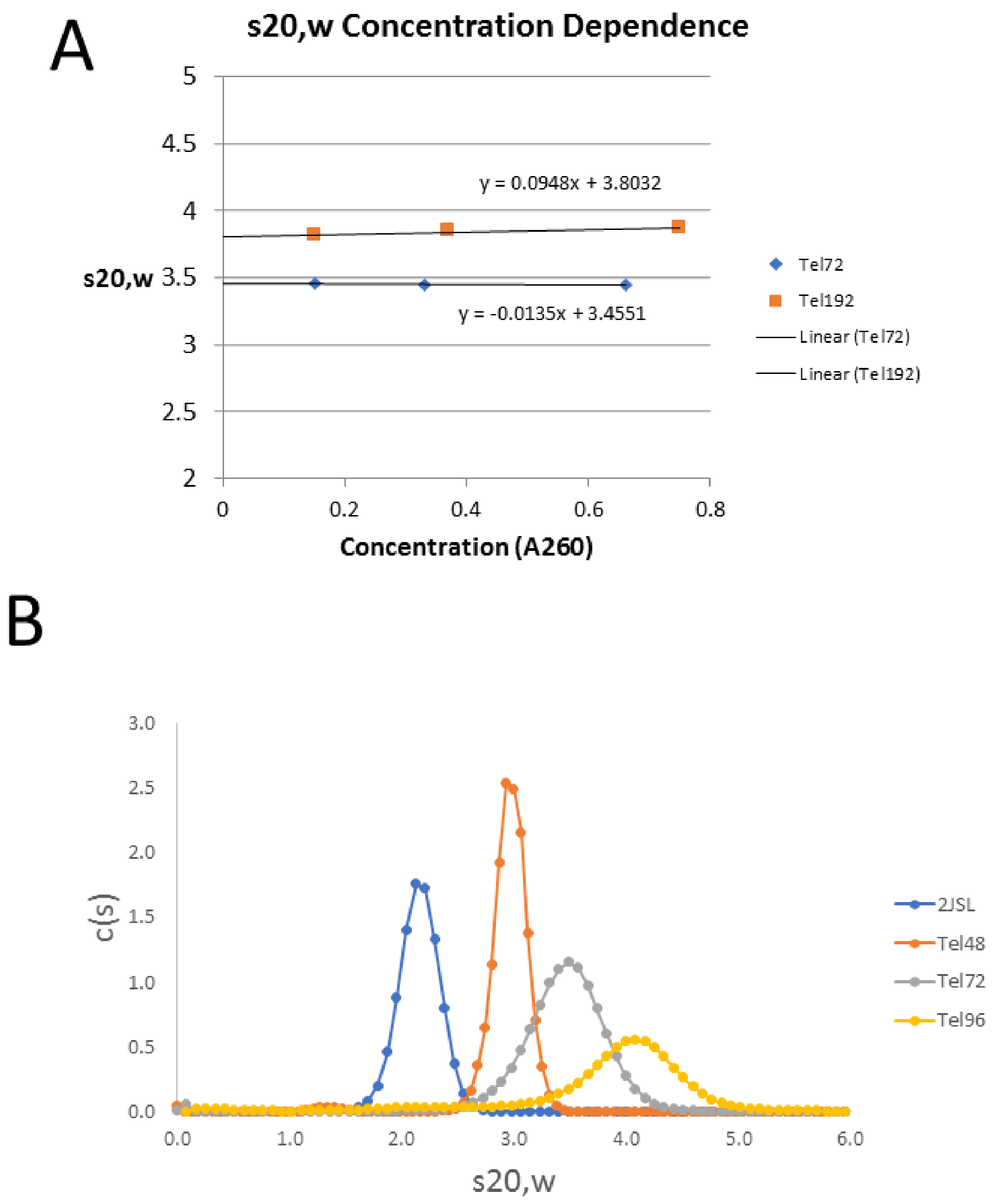

**Figure S2.** Sedimentation velocity analysis of higher-order telomere sequences. (Top) Analysis of Tel72 and Tel96 concentration dependence on sedimentation. Extrapolation to the Y-axis gives the infinite dilution $S_{20,w}$ values. (Bottom) representative C(s) vs. $S_{20,w}$ distributions for 2JSL, Tel48, Tel72, and Tel96.

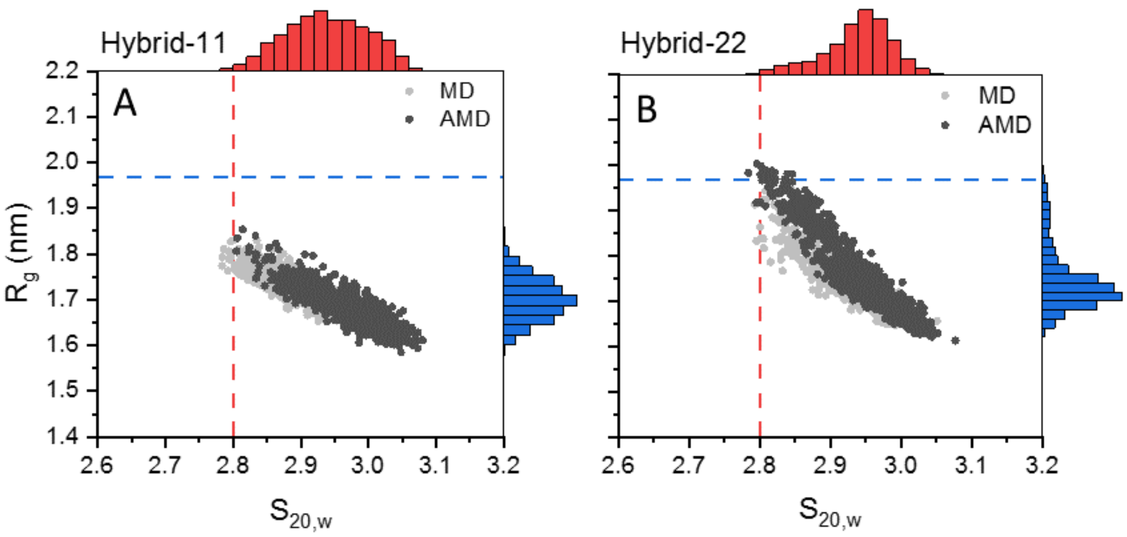

**Figure S3.** Additional results of Tel48 SAXS atomistic modeling efforts shown in **Figure 2** of main text. (A-B) scatter plots of calculated radii of gyration and sedimentation coefficients for hybrid-11 (A) and hybrid-22 (B) with MD-derived values shown in light gray and aMD-derived values in dark gray. The inset dashed red and blue lines represent the experimentally measured values for sedimentation coefficient and radius of gyration, respectively. The outer histograms represent the distributions of values from both MD and aMD snapshots combined. The histograms indicate that the major sampled conformations in both cases are much more compact than would be expected from either SAXS or AUC analyses.

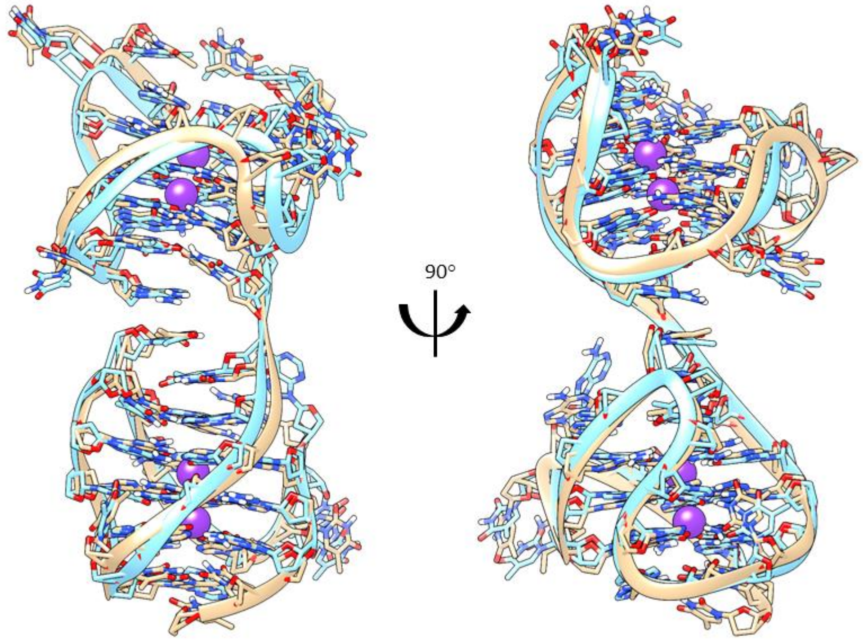

**Figure S4.** Comparison of the Tel48 (cyan) and Tel50 (tan) hybrid-12 conformers. The tan hybrid-12 conformer was taken from an earlier report by Petraccone et al.(1) and the cyan hybrid-12 is the model derived here from EOM (bottom right-most conformer in **Figure 2D**). The pair-wise residue RSMD is 1.6 Å as determined by the matchmaker module of UCSF Chimera v1.12. Potassium is shown as purple spheres and is derived from the Petraccone model (EOM hybrid-12 potassium is hidden).

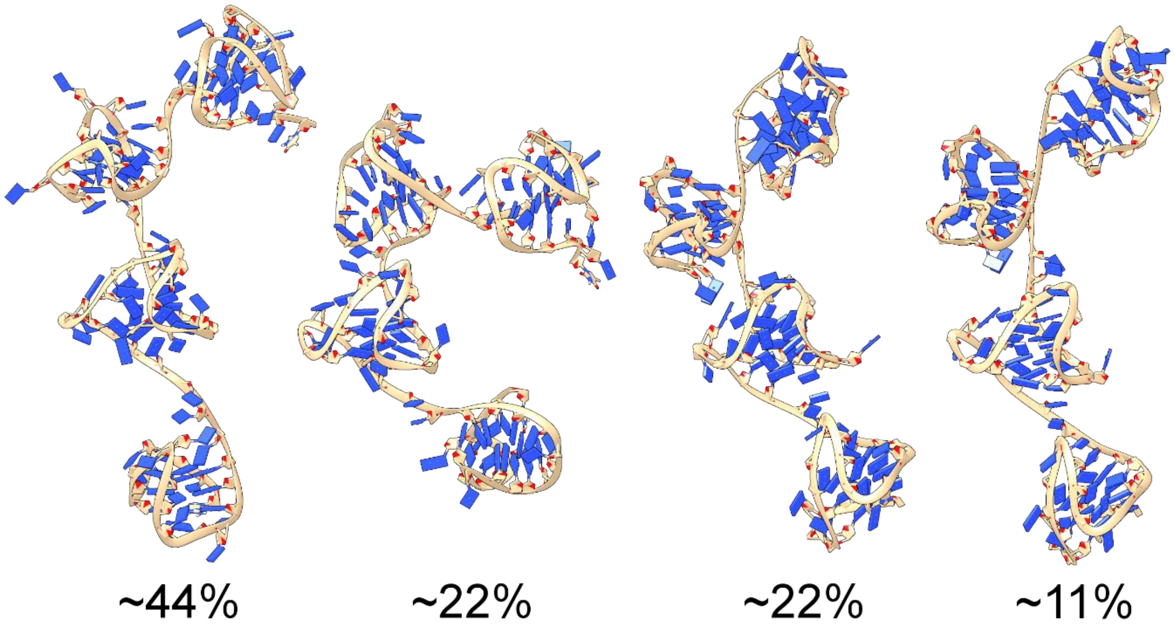

**Figure S5.** Results of Tel96 atomistic modeling efforts. Depicted are the four hybrid-2122 conformers derived from EOM analysis of the 100 ns MD simulations with their respective weights (as % of the reconstructed scattering curve). All conformers are arranged with their 5' ends at the top of the figure.

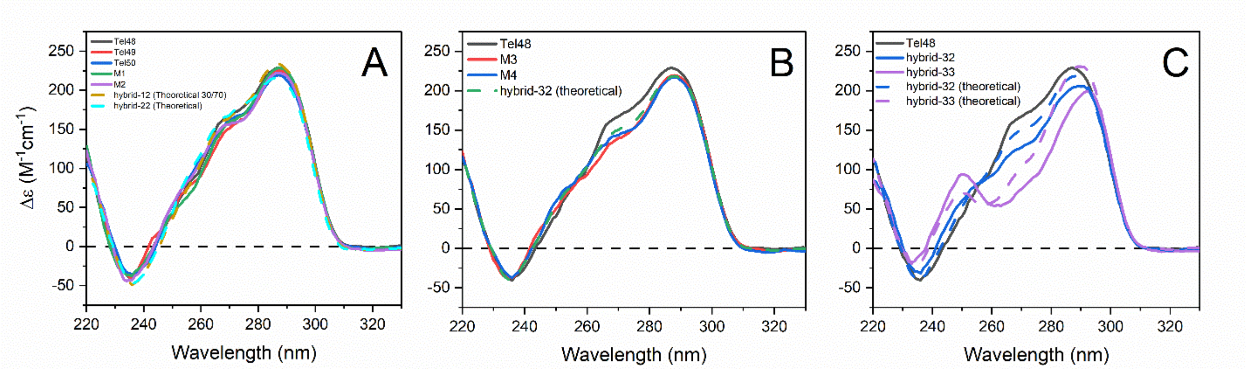

**Figure S6.** Normalized CD spectra of Tel48 compared to various flanking residue and internal mutant sequences.

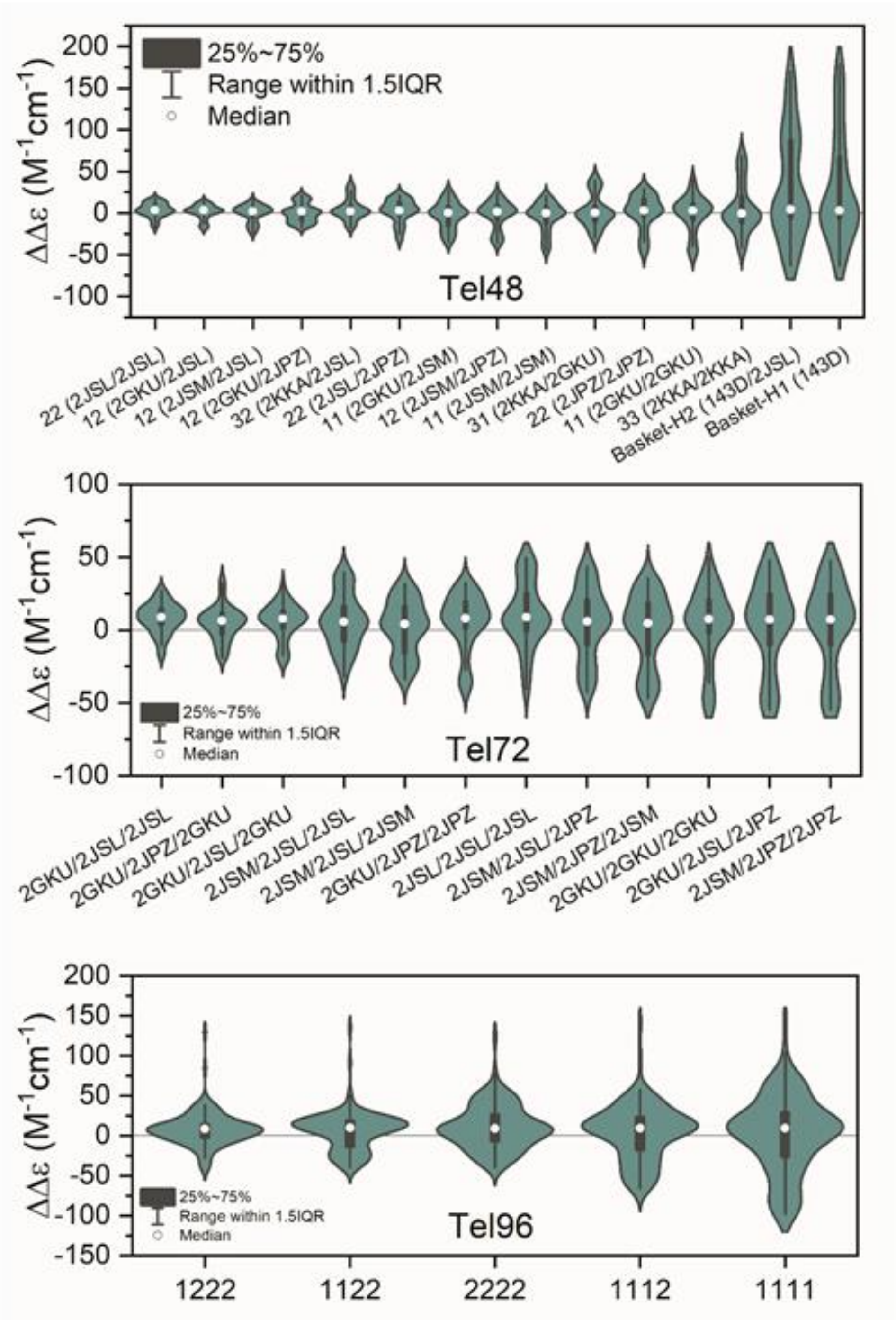

**Figure S7.** Violin plots of residuals obtained as the difference between experimental and theoretical CD reconstruction curves for Tel48, Tel72, and Tel96. The residuals are plotted from best (left) to worst (right) based on residual sum of squares analysis in Origin 2020.
